# TNF receptor associated factor 6 interacts with ALS-linked misfolded superoxide dismutase 1 and promotes aggregation

**DOI:** 10.1101/780460

**Authors:** Sabrina Semmler, Myriam Gagné, Pranav Garg, Sarah Pickles, Charlotte Baudouin, Emeline Hamon-Keromen, Laurie Destroismaisons, Yousra Khalfallah, Mathilde Chaineau, Elise Caron, Andrew N. Bayne, Jean-François Trempe, Neil R. Cashman, Alexandra T. Star, Arsalan S. Haqqani, Thomas M. Durcan, Elizabeth M. Meiering, Janice Robertson, Nathalie Grandvaux, Steven S. Plotkin, Heidi M. McBride, Christine Vande Velde

## Abstract

Amyotrophic lateral sclerosis (ALS) is a fatal disease, characterized by the selective loss of motor neurons leading to paralysis. Mutations in the gene encoding superoxide dismutase *1* (SOD1) are the second most common cause of familial ALS, and considerable evidence suggests that these mutations result in an increase in toxicity due to protein misfolding. We previously demonstrated in the SOD1^G93A^ rat model that misfolded SOD1 exists as distinct conformers and forms deposits on mitochondrial subpopulations. Here, using SOD1^G93A^ rats and conformation-restricted antibodies specific for misfolded SOD1 (B8H10 and AMF7-63), we identified the interactomes of the mitochondrial pools of misfolded SOD1. This strategy identified binding proteins that uniquely interacted with either AMF7-63 or B8H10-reactive SOD1 conformers as well as with a high proportion of interactors common to both conformers. Of this latter set, we identified the E3 ubiquitin ligase TNF receptor-associated factor 6 (TRAF6) as a SOD1 interactor and determined that exposure of the SOD1 functional loops facilitates this interaction. Of note, this conformational change was not universally fulfilled by all SOD1 variants and differentiated TRAF6-interacting from TRAF6 non-interacting SOD1 variants. Functionally, TRAF6 stimulated polyubiquitination and aggregation of the interacting SOD1 variants. TRAF6 E3 ubiquitin ligase activity was required for the former, but was dispensable for the latter, indicating that TRAF6-mediated polyubiquitination and aggregation of the SOD1 variants are independent events. We propose that the interaction between misfolded SOD1 and TRAF6 may be relevant to the etiology of ALS.

---

Amyotrophic lateral sclerosis (ALS) is a progressive paralytic and ultimately fatal disease, characterized by the degeneration of upper and lower motor neurons in the brain, brainstem and spinal cord. Mutations in the *SOD1* gene, encoding superoxide dismutase 1, were the first to be identified as causative of ALS (1). *SOD1* mutations are the second most common genetic cause of ALS to date, accounting for about 12% of familial cases (fALS) and 1% of sporadic ALS cases (sALS) (2). Over 180 different mutations in *SOD1* have thus far been reported as pathogenic (<u>alsod.iop.kcl.ac.uk</u>). These mostly comprise missense mutations in coding regions, but rare nonsense mutations, deletions, insertions, and mutations in non-coding regions have also been reported (3). Mutations in the coding region are found in all five exons of *SOD1*. It is now agreed that SOD1-mediated ALS rises from a toxic gain-of-function due to mutation-induced changes in the folding of the SOD1 protein, collectively referred to as misfolded SOD1.

Misfolded SOD1 exposes certain structural features, normally buried in natively folded SOD1, that have been exploited for the generation of antibodies that detect misfolded SOD1 on a conformation-restricted basis (4,5). Amongst those, the mouse monoclonal B8H10 recognizes an epitope within the metal-binding loop (loop IV, within aa. 57-80), the exposure of which is associated with a deficiency in metal co-factor binding (6–8). The rabbit monoclonal AMF7-63 is the high-affinity descendant of the mouse monoclonal DSE2 (disease-specific epitope) and was raised against residues in the electrostatic loop (loop VII, aa. 125-142), the exposure of which is associated with increased conformational flexibility in this region (9,10). Strategies to target and thus eliminate misfolded SOD1 in transgenic rodent models, using either active or passive immunization approaches, show benefit in delaying disease phenotypes (6,8,11–15). Whether or not a common target exists for misfolded SOD1 remains unknown. Using B8H10 and AMF7-63, we have previously demonstrated in the SOD1^G93A^ rat model of ALS that different conformers of misfolded SOD1 deposit on the surface of spinal cord mitochondria in an age-dependent manner, supporting the view that distinct misfolded SOD1 conformers exist (10,16). Interestingly, the accumulation of both conformers positively correlates with mitochondrial damage (10,16,17).

Here, we report on the identification of the E3 ubiquitin ligase TNF receptor-associated factor 6 (TRAF6) as a novel binding partner of misfolded SOD1. In cellulo, we show that the interaction with TRAF6 is unique to mutant but not wild type SOD1 and we map the interaction to the C-terminus of TRAF6. However, not all SOD1 variants interact equally with TRAF6. We show that TRAF6-interacting and non-interacting SOD1 variants are conformationally distinct in terms of their exposure of misfolding-associated epitopes recognized by B8H10 and AMF7-63. We demonstrate that TRAF6 stimulates mutant SOD1 polyubiquitination and aggregation, but that these variably depend on TRAF6 E3 ubiquitin ligase activity and are therefore independent events. Specifically, siRNA-mediated reduction of TRAF6 levels reduces mutant SOD1 aggregation in a cellular model. Importantly, we uncover that TRAF6 localization is not restricted to the cytoplasm, with a pool of TRAF6 being constitutively localized to spinal cord mitochondria. Lastly, TRAF6 is expressed in the cell types harboring misfolded SOD1 *in vivo*, establishing relevance to misfolded SOD1-linked pathology in ALS. Taken together, these results suggest that misfolded SOD1 interacts with a mitochondrial-localized pool of TRAF6 and this interaction may underlie misfolded SOD1 accumulation.

## RESULTS

### Identification of novel binding partners of mitochondria-associated misfolded SOD1

In mutant SOD1 rodent models of ALS, a portion of misfolded SOD1 conformers are localized to the surface of spinal cord mitochondria (10,16–18). To identify the interactomes of specific misfolded SOD1 conformers at the mitochondria, we performed immunoprecipitation coupled with mass spectrometry (IP-MS) of spinal cord mitochondria isolated from symptomatic SOD1^G93A^ rats (**Fig. 1A**). Briefly, misfolded SOD1 was immunoprecipitated with anti-B8H10 or anti-AMF7-63, and eluates were analyzed by mass spectrometry. As previously published, both B8H10 and AMF7-63 specifically detect misfolded forms of SOD1 but are unreactive for natively-folded SOD1^WT^ (8–10,16,17). From all identified prey proteins, only those with a maximum peptide score ≥40 in three of three biological replicates and not detected in IgG controls were considered as potential binding partners. Identified proteins are listed in **Table S1**. Human SOD1 was detected in all samples and served as an internal control. Also, known binding partners of SOD1 [e.g. the copper chaperone for SOD1 (CCS) (19)] and misfolded SOD1 [e.g. voltage-gated anion channel 1 (VDAC1) (20)] were detected, which additionally supports the quality and validity of our dataset. Of the 52 proteins identified, 3 proteins (6%) interacted uniquely with B8H10-reactive SOD1 conformers and 7 uniquely interacted with AMF7-63 reactive conformers (13%) (**Fig. 1B**, **Table S2**). Thus, the majority of interactors (81%) were common to B8H10- and AMF7-63-reactive conformers of misfolded SOD1.

**Figure 1:**
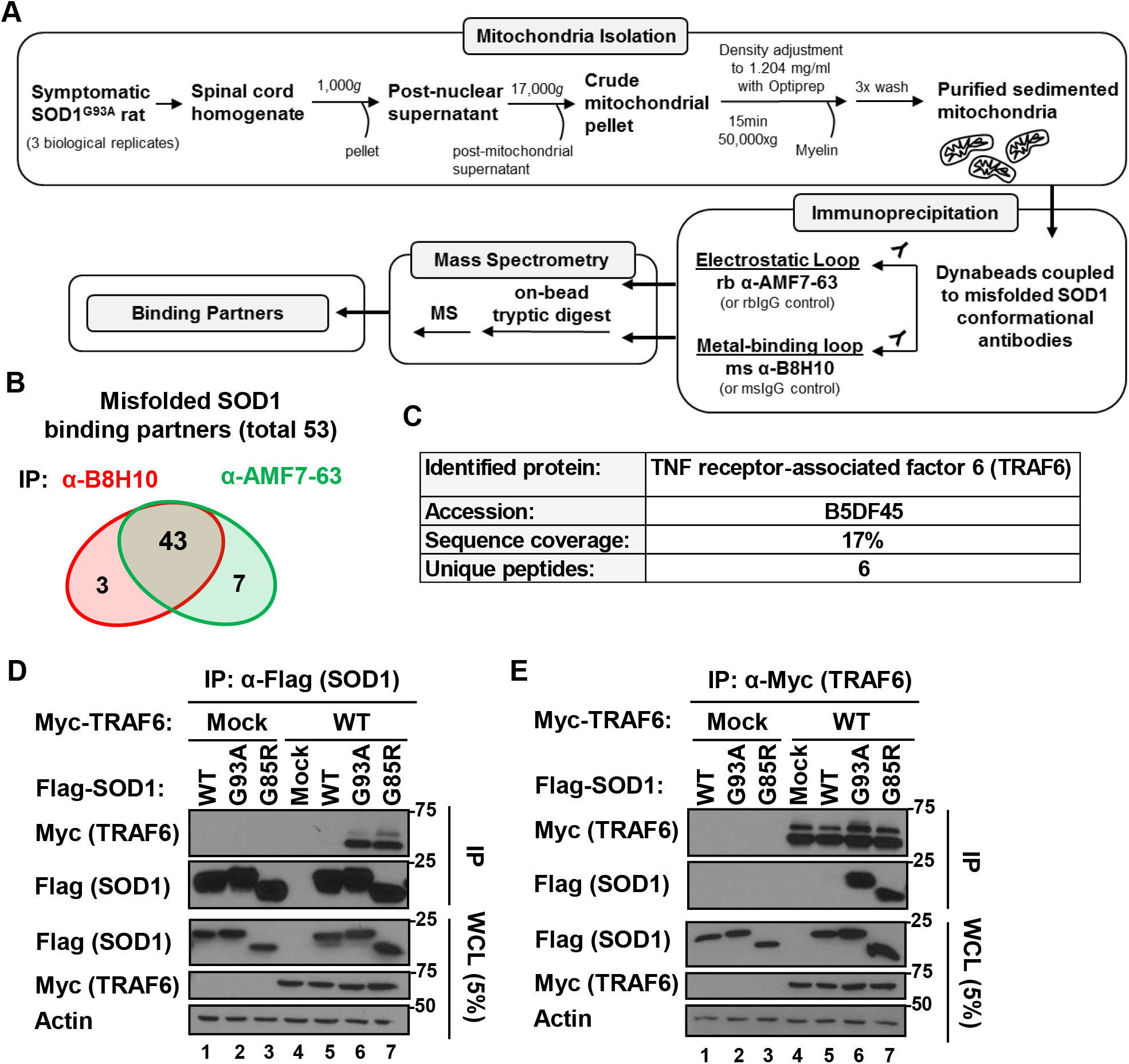
TRAF6 is a novel binding partner of B8H10 and AMF7-63-reactive misfolded SOD1 at the mitochondria. **(A)** Workflow for immunoprecipitation-based mass spectrometry proteomics for the identification of binding partners of mitochondria-associated misfolded SOD1 in the SOD1^G93A^ rat model. Misfolded SOD1 was immunoprecipitated with the misfolded SOD1-specific conformation-restricted antibodies B8H10 and AMF7-63 from SOD1^G93A^ rat spinal cord mitochondria and binding partners identified by nanoLC-MS/MS. **(B)** Venn diagram of identified misfolded SOD1 binding partners common to both B8H10 and AMF7-63 conformers, or selective for either. **(C)** The novel binding partner TRAF6 (UniProt ID: B5DF45) was identified based on 6 unique peptides covering 17% of the protein sequence. **(D, E)** Flag-tagged SOD1 (wild type or mutants) and Myc-tagged TRAF6 (wild type or deletion mutants) were co-expressed in 293FT cells. Co-immunoprecipitations were performed on Flag-SOD1, or reciprocally on Myc-TRAF6, and co-precipitation of either binding partner was analyzed by immunoblotting. Immunoblotting for Flag-SOD1 (bait) is to demonstrate equal IP efficiency across conditions. Mock refers to transfection with an equivalent amount of empty vector. Note, the slower migrating band in TRAF6 lanes is auto-ubiquitinated TRAF6. Whole cell lysates (WCL) were loaded to demonstrate equal plasmid expression. Actin serves as loading control. Data is representative of at least 3 independent experiments.

To extract information on the biological significance of our dataset, we used Enrichr to perform functional annotation and pathway enrichment analyses (21,22). Our dataset revealed a significant enrichment for molecules participating in biological processes, such as *response to unfolded protein* (GO:006986, q-value = 1.47E^-04^), *removal of superoxide radicals* (GO:0019430, q-value = 5.66E^-04^), *succinyl-CoA metabolic process* (GO:0006104, q-value = 3.12E^-^ 02), *intermediate filament bundle assembly* (GO:0045110, q-value = 2.84E^-02^), and *cellular response to interleukin-12* (GO: 0071349, q-value = 4.44E^-02^) (**Table S3**). On the basis of molecular function, our dataset significantly accumulated molecules that are *C3HC4-type RING finger domain binding* (GO:0055131, q-value = 3.52E^-^ 04), or possess *ATPase activity* (GO:0016887, q-value = 4.87E^-02^) (**Table S3**). Next, KEGG pathway analysis showed a significant enrichment for molecular players associated not only with *Amyotrophic lateral sclerosis* (q-value = 1.90E^-05^), but also *Huntington disease* (q-value = 3.91E^-04^), *Parkinson disease* (q-value = 1.24E^-02^), and *Alzheimer disease* (q-value = 2.10E^-02^). In addition, our dataset revealed an accumulation of molecules participating in *antigen processing and presentation* (q-value = 2.32E^-02^) and infectious disease pathways (**Table S4**).

### Presence of ubiquitin in AMF7-63 and B8H10 immunoprecipitates

Ubiquitin is highly conserved and encoded by four different genes: *UBB*, *UBC*, *UBA52* and *RPS27A*. While *UBB* and *UBC* encode polyubiquitin precursors made up of multiple head-to-tail repeats of ubiquitin, *UBA52* and *RPS27A* are ubiquitin hybrid genes encoding a fusion protein comprised of a single N-terminal ubiquitin moiety fused to a C-terminal extension. In reviewing our misfolded SOD1 interactomes, we noticed that ubiquitin-60S ribosomal protein L40 (RL40), which is encoded by *UBA52*, was a binding partner detected in both B8H10 and AMF7-63 immunoprecipitates (**Table S1**, **Fig. S1A**). To elucidate whether the peptides listed under RL40 correspond to the N-terminal ubiquitin moiety or the C-terminal extension of the protein, we generated a sequence alignment for rat RL40 with ubiquitin (single moiety) (**Fig. S1B**). All three peptides listed under the identifier RL40 mapped to the N-terminal ubiquitin moiety and not the C-terminal extension of the protein. This indicates that the detected peptides are not unique for RL40, but generally correspond to ubiquitin and may thus derive from RL40, RS27A, UBB or UBC proteins. To illustrate this, we also aligned rat RS27A (**Fig. S1B**). To investigate whether ubiquitin is indeed associated with B8H10- and AMF7-63-reactive misfolded SOD1, we precipitated misfolded SOD1 from SOD1^G93A^ rat spinal mitochondria and immunoblotted for ubiquitin (**Fig. S1C**). In both B8H10- and AMF7-63-immunoprecipitates, ubiquitin was detected as a high molecular weight smear refusing entry into the separating gel, a pattern typically observed with intensely polyubiquitin-modified proteins. Taken together, these data suggest that B8H10 and AMF7-63 conformers are modified with ubiquitin (or polyubiquitin) or associate with stable binding partners that are themselves ubiquitinated.

### Identification of the E3 ubiquitin ligase TRAF6 as a novel binding partner of misfolded SOD1

In our dataset, we detected six unique peptides for an E3 ubiquitin ligase: TNF receptor-associated factor 6 (TRAF6). TRAF6 peptides, covering 17% of the protein sequence, were detected in both B8H10 and AMF7-63 immunoprecipitates (**Table S1**, **Fig. 1C**, **Fig. S2**).

TRAF6 is a cytoplasmic-localized RING-type ubiquitin ligase classically known for its role in the canonical nuclear factor kappa-B (NF-κB) signaling pathway, where it participates in the transduction of signals downstream of a multitude of immunoregulatory receptors (23–26) and functions as a hub protein in the crosstalk of immune-regulatory pathways with the autophagy machinery (27–30). TRAF6 is not strictly localized to the cytoplasm but translocates to mitochondria in certain contexts (31–35). Importantly, TRAF6 has been previously implicated in neurodegenerative diseases, including Parkinson’s disease (36–38), Huntington’s disease (39), and Alzheimer’s disease (40–42), where it stimulates the ubiquitination of disease-associated proteins and exerts a regulatory role on their turnover and aggregation (43). The detection of TRAF6 as a binding partner of misfolded SOD1 was additionally intriguing, since physiological interactions between E3 ubiquitin ligases and their substrates are generally known to be weak, transient, and not readily detected by IP-MS based techniques (44–46). We speculated that the interaction is unnaturally stable and of possible pathological relevance. Thus, we elected to more fully investigate the interaction between mutant/misfolded SOD1 and TRAF6.

### TRAF6 interacts with mutant but not wild type SOD1

To characterize the interaction between mutant SOD1 and TRAF6, we established an in cellulo system based on transient co-expression of Flag-tagged SOD1 and Myc-tagged TRAF6 in human 293FT cells. First, to elucidate whether this system can recapitulate our *in vivo* results, we performed reciprocal co-immunoprecipitations (**Fig. 1D, E**). SOD1^G93A^ and SOD1^G85R^, but not SOD1^WT^, co-precipitated TRAF6^WT^ (**Fig. 1D**, lanes 5-7). Reversely, TRAF6^WT^ co-precipitated mutant SOD1, but not SOD1^WT^ (**Fig. 1E**, lanes 5-7), thus confirming our IP-MS results. Note, the slower-migrating TRAF6 band is the auto-ubiquitinated form, consistent with previous work (47). In addition, these data indicate that TRAF6 selectively interacts with mutant but not wild type SOD1, indicating that the interaction with TRAF6 is unique to non-native SOD1 conformers.

### Mutant SOD1 binds the TRAF6 C-terminus

TRAF6 is a multidomain protein and highly conserved across mammalian species (48,49). The N-terminus features a really interesting new gene domain (RING) with a series of zinc finger motifs (zF), which are connected via an α-helical coiled-coil motif (CC) to a C-terminal meprin and TRAF homology (MATH) domain (**Fig. 2A, B**). The RING domain and adjacent zinc finger motifs are involved in E2 ubiquitin conjugase binding and are required for TRAF6 function as an E3 ubiquitin ligase (50). The MATH domain is an eight-stranded anti-parallel β-sandwich with which TRAF6 engages in a multitude of protein-protein interactions (32,51–53). To determine the domain of TRAF6 that interacts with mutant SOD1, two TRAF6 deletion constructs were generated: TRAF6^ΔC^ (aa. 1-288) comprising the N-terminal RING domain and zinc finger motifs, and TRAF6^ΔN^ (aa. 289-522) comprising the central coiled-coil motif and C-terminal MATH domain (**Fig. 2A, B**). SOD1^G93A^ and SOD1^G85R^ efficiently co-precipitated TRAF6^ΔN^, but not TRAF6^ΔC^ (**Fig. 2C**, compare lanes 14-15 to lanes 11-12). SOD1^WT^ did not interact with TRAF6^WT^, nor the two TRAF6 deletion mutants (**Fig. 2C**, lanes 7, 10 and 13). We conclude that the C-terminus of TRAF6 is necessary and sufficient for mutant SOD1 binding.

**Figure 2:**
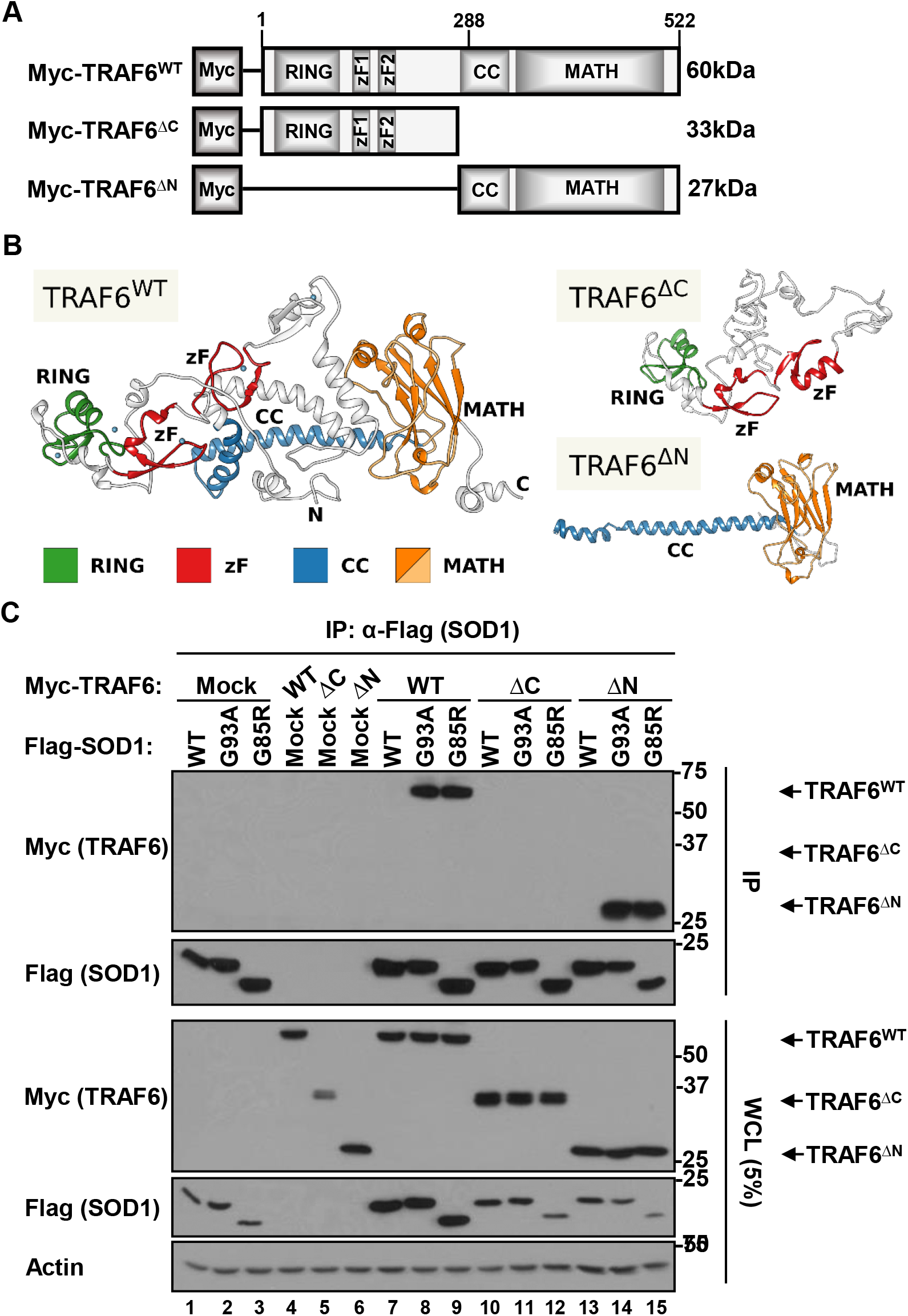
The C-terminus of TRAF6 is necessary and sufficient for mutant SOD1 binding. **(A)** Schematic representation of generated Myc-TRAF6 deletion constructs. TRAF6^ΔC^ (aa. 1-288) comprises the N-terminal RING domain and zinc finger motifs. TRAF6^ΔN^ (aa. 289-522) comprises the central coiled-coil motif (CC) and C-terminal MATH domain. **(B)** Protein structure models for TRAF6^WT^, TRAF6^ΔC^, and TRAF6^ΔN^. **(C)** Flag-tagged SOD1 (wild type or mutants) and Myc-tagged TRAF6 (wild type or deletion mutants) were co-expressed in 293FT cells. Co-immunoprecipitations were performed on Flag-SOD1 and Myc-TRAF6 co-precipitation was assessed by immunoblotting. Immunoblotting for Flag-SOD1 (bait) is to demonstrate equal IP efficiency across conditions. Mock refers to transfection with an equivalent amount of empty vector. Whole cell lysates (WCL) were loaded to demonstrate equal plasmid expression. Actin serves as loading control. Data is representative of at least 3 independent experiments.

### SOD1 mutation-dependent variation in TRAF6 binding

To evaluate if the interaction of mutant SOD1 with TRAF6 is a general property of SOD1 variants, we generated Flag-tagged expression plasmids for seven additional fALS-associated SOD1 missense mutations: SOD1^A4V^, SOD1^G37R^, SOD1^H46R^, SOD1^E100G^, SOD1^V148G^, and SOD1^V148I^. We also cloned SOD1^G127X^, a fALS-associated SOD1 truncation mutant lacking the C-terminus. Again, we co-expressed these mutants with Myc-TRAF6 (wild type or ΔN) in 293FT cells, immunoprecipitated Flag-SOD1 from cell lysates, and assessed Myc-TRAF6 co-precipitation by immunoblot (**Fig. 3A, B**). As previously demonstrated, SOD1^G93A^ and SOD1^G85R^ co-precipitated TRAF6^WT^ to equivalent amounts (**Fig. 1D, E**). A similar level of interaction was also observed between TRAF6^WT^ and SOD1^E100G^, SOD1^G127X^, SOD1^V148G^, and SOD1^G37R^, while SOD1^A4V^ showed the most robust interaction (**Fig. 3A**, lanes 7-13). In contrast, SOD1^H46R^ very weakly co-precipitated TRAF6^WT^ (**Fig. 3A**, lane 6). Interestingly, SOD1^V148I^ did not co-precipitate with TRAF6^WT^ (**Fig. 3A**, lane 5), nor TRAF6^ΔN^ (**Fig. 3C**, lane 11). Taken together, there is variation in TRAF6 binding by SOD1 mutant proteins.

**Figure 3:**
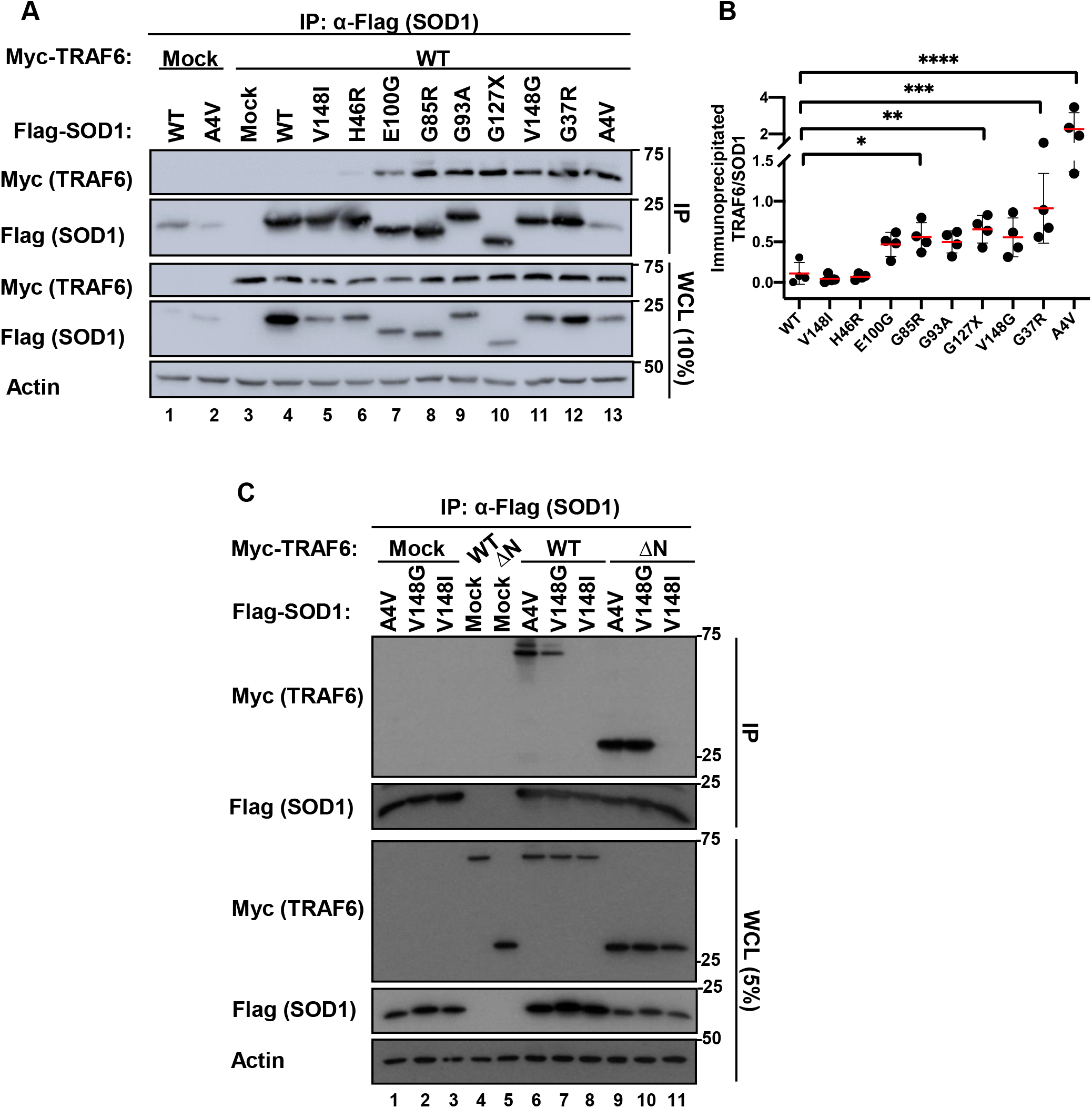
SOD1 mutation-dependent variation in the interaction with TRAF6 in cellulo. **(A-C)** Flag-tagged SOD1 (wild type or mutants) and Myc-tagged TRAF6 (wild type or ΔN) were co-expressed in 293FT cells. Co-immunoprecipitations were performed on Flag-SOD1 and Myc-TRAF6 co-precipitation analyzed by immunoblot. Immunoprecipitates were immunoblotted for Flag-SOD1 to demonstrate equal IP efficiency across conditions. Mock refers to transfection with an equivalent amount of empty vector. Whole cell lysates (WCL) were loaded to demonstrate equal plasmid expression. Actin serves as loading control. **(B)** Densitometric analysis of the co-immunoprecipitated amount of Myc-TRAF6 relative to the amount of immunoprecipiated Flag-SOD1. Plotted values are the mean ± SD of 4 independent experiments. * p<0.05, ** p<0.01, *** p<0.001, **** p<0.0001.

Consistent with the view that more than one misfolded SOD1 conformer exists (10,54), these data suggest that many SOD1 mutant proteins adopt a misfolded conformation(s) that is (are) favorable for TRAF6 binding, but that these are not conformationally identical, which may account for the observed variable interaction with TRAF6. Interestingly, one ALS-causing mutant, SOD1^V148I^, did not interact with TRAF6, consistent with previous data indicating that this mutant is WT-like in terms of stability and structure (55–57). Strikingly, the substitution at Val-148 with glycine is sufficient for mutant SOD1 to adopt a TRAF6-interacting conformation.

Lastly, the variation in mutant SOD1 binding was observed for TRAF6^WT^ but not for TRAF6^ΔN^. SOD1^A4V^ and SOD1^V148G^ co-precipitated TRAF6^WT^ to variable amounts but TRAF6^ΔN^ to similar amounts (**Fig. 3C**, compare lanes 6-7 to lanes 9-10). While the TRAF6 C-terminus is required for mutant SOD1 binding, these data suggest that the N-terminus of TRAF6 modulates the interaction with mutant SOD1.

### SOD1 misfolding-associated functional loop exposure is required for TRAF6 interaction

Next, we sought to investigate the relevance of SOD1 misfolding-associated exposure of the metal-binding loop (loop IV) and electrostatic loop (loop VII) to TRAF6 binding. For this, Flag-SOD1 variants expressed in 293FT cells were evaluated for reactivity to the conformation-restricted misfolded SOD1 antibodies B8H10 (epitope in loop IV) or AMF7-63 (epitope in loop VII) by immunoprecipitation (**Fig. 4A, B**). We incorporated the C-terminal truncation mutant SOD1^G127X^ as a negative control for AMF7-63 immunoprecipitations, as this mutant lacks the electrostatic loop (loop VII). While most SOD1 variants immunoprecipitated with both B8H10 and AMF7-63, SOD1^V148I^ was not immunoprecipitated with either antibody (**Fig. 4B**, compare lanes 3-9 to lane 10). SOD1^G127X^ was not immunoprecipitated with AMF7-63 (as expected), but was with B8H10 (**Fig. 4B**, lane 11). As expected, neither antibody immunoprecipitated SOD1^WT^ (**Fig. 4B**, lane 2). To summarize, all TRAF6-interacting SOD1 variants were reactive for either B8H10 and/or AMF7-63. In contrast, SOD1^WT^ and SOD1^V148I^, which both do not interact with TRAF6, were unreactive for both antibodies suggesting that SOD1 misfolding-associated exposure of the functional loops or monomerization/unfolding of SOD1 are conformational requirements for TRAF6 interaction. Lastly, data obtained with SOD1^G127X^ indicates that exposure of the metal-binding loop is sufficient for TRAF6 binding.

**Figure 4:**
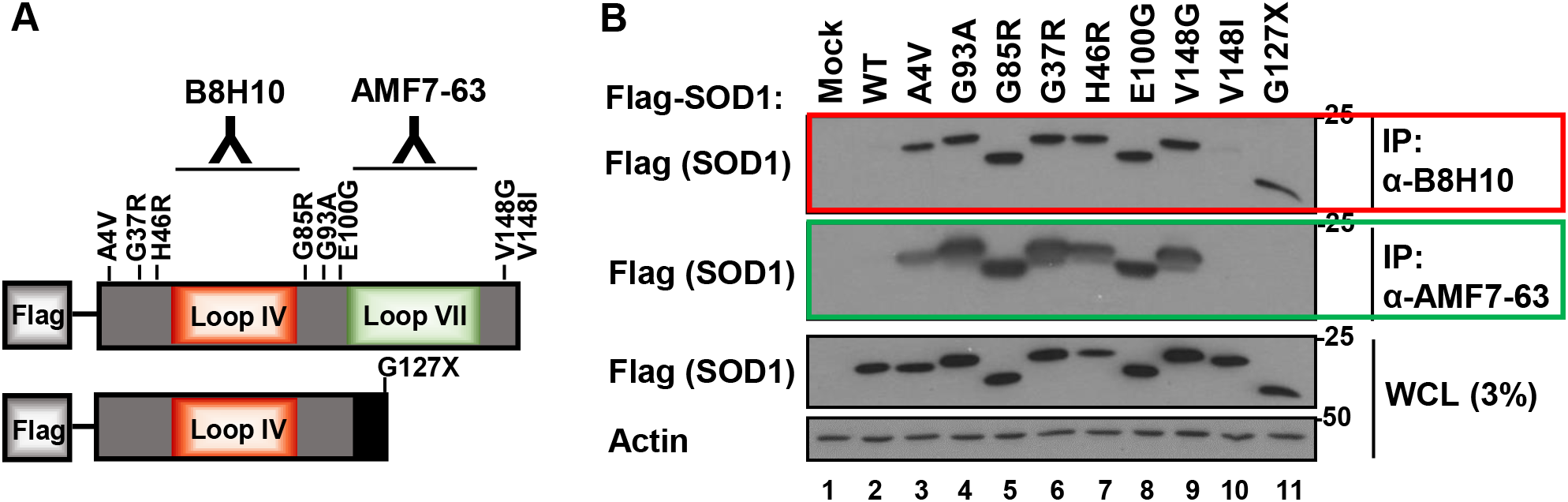
Exposure of epitopes associated with SOD1 misfolding discerns TRAF6-interacting from non-interacting mutants. **(A)** Schematic representation of Flag-SOD1 constructs, with the location of mutation sites relative to B8H10 (loop IV) and AMF7-63 (loop VII) antibody epitopes. SOD1^G127X^ is a truncation mutant with a non-native C-terminus and inherently lacks the AMF7-63 epitope. **(B)** Flag-tagged SOD1 (wild type or mutants) was expressed in 293FT cells. Misfolded SOD1 was immunoprecipitated with the B8H10 or AMF7-63 antibody and precipitates were analyzed by western blot. Mock refers to transfection with an equivalent amount of empty vector. Whole cell lysates (WCL) were loaded to demonstrate equal plasmid expression. Actin serves as loading control. Data is representative of 3 independent experiments.

### TRAF6 stimulates mutant SOD1 ubiquitination

TRAF6 is reported to stimulate the ubiquitination and aggregation of mutant/misfolded proteins in the context of other neurodegenerative diseases (36,39,42). To determine if mutant SOD1 is also a substrate of TRAF6, we performed in cellulo ubiquitination assays (**Fig. 5A**). For this, we co-expressed Flag-SOD1 and Myc-TRAF6^WT^ or E3 ubiquitin ligase-inactive mutant TRAF6^C70A^ (58) with HA-Ubiquitin, immunoprecipitated HA-Ubiquitin from whole cell lysates and immunoblotted for Flag-SOD1. In this assay, we selected an SOD1 variant which demonstrates robust interaction with TRAF6 and is dually reactive for B8H10 and AMF7-63 (SOD1^A4V^), an SOD1 variant that exhibits a slightly less robust in interaction with TRAF6 and also reactive for B8H10 and AMF7-63) (SOD1^V148G^) and a variant that does not interact with TRAF6 and does not display B8H10 or AMF7-63 reactive SOD1 epitopes (SOD1^V148I^). TRAF6 strongly stimulated the polyubiquitination of SOD1^A4V^ and SOD1^V148G^, but not SOD1^V148I^ (**Fig. 5A**, compare lanes 5-7). The polyubiquitin signature on SOD1^A4V^ and SOD1^V148G^ was absent in cells co-expressing TRAF6^C70A^ (**Fig. 5A**, compare lanes 5-6 to 8-9). These data indicate that TRAF6 can stimulate the polyubiquitination of mutant SOD1 in an E3 ubiquitin ligase activity-dependent manner. Moreover, this effect is selective to TRAF6-interacting SOD1 variants and the level of detected polyubiquitin-modified mutant SOD1 positively correlates with the observed differences in TRAF6-mutant SOD1 interaction.

**Figure 5:**
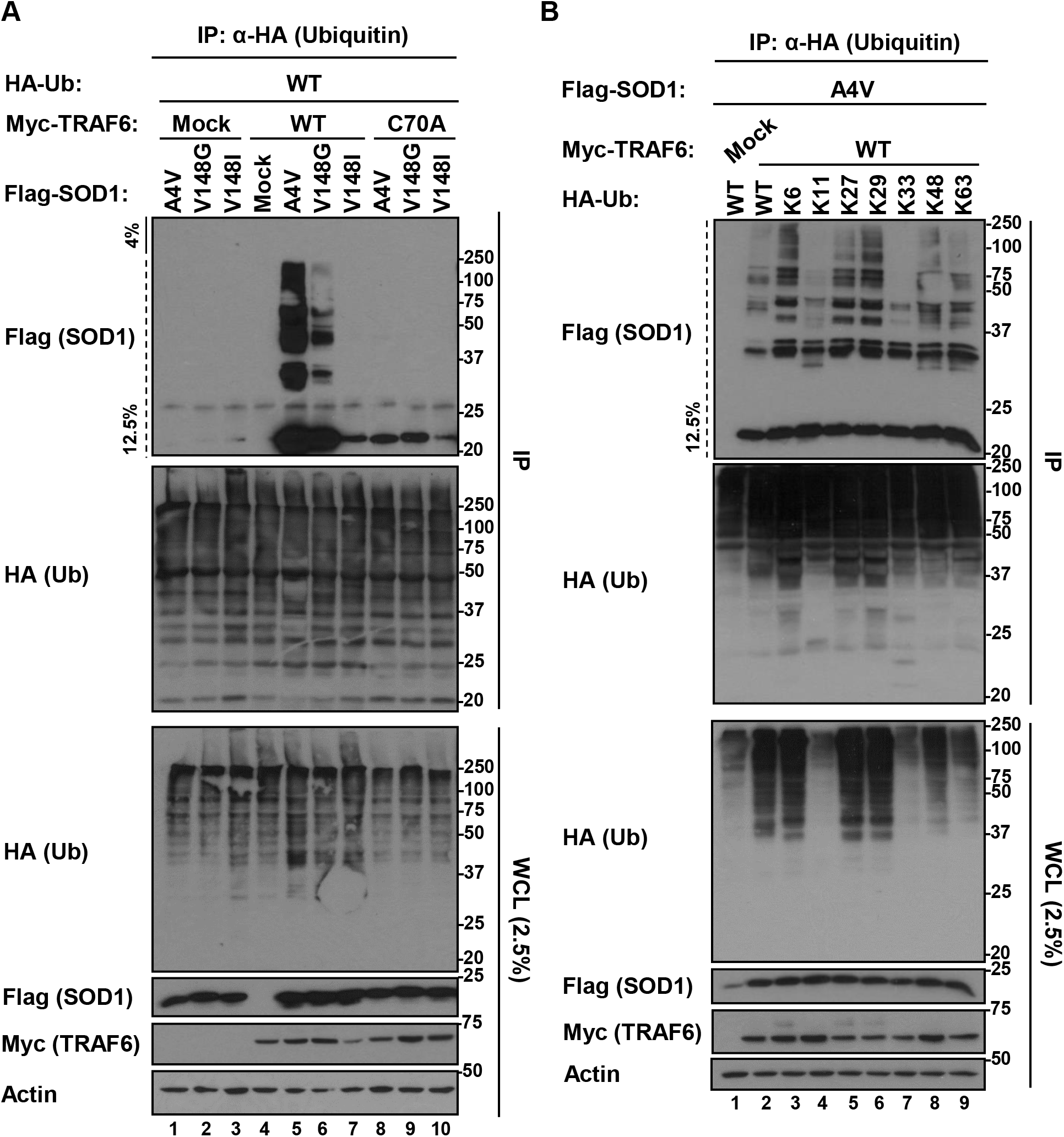
TRAF6-mediated polyubiquitination of mutant SOD1 depends on its ubiquitin ligase activity and is selective to TRAF6-interacting SOD1 variants. **(A)** In cellulo ubiquitination assays with mutant SOD1 as substrate and TRAF6 as catalyzing E3 ubiquitin ligase. Flag-SOD1^A4V^ and SOD1^V148G^ (TRAF6-interacting), or Flag-SOD1^V148I^ (TRAF6 non-interacting), were co-expressed with Myc-TRAF6^WT^ or ubiquitin ligase-inactive TRAF6^C70A^, in the presence of HA-Ubiquitin^WT^ in 293FT cells. Immunoprecipitations were performed on HA-Ubiquitin and immunoprecipitates were immunoblotted for Flag-SOD1, and HA-Ubiquitin to demonstrate equal IP efficiency across conditions. **(B)** In cellulo ubiquitin linkage assays were performed akin to panel A, but in the presence of HA-Ubiquitin^WT^ or single-lysine mutants. TRAF6 mediates mutant SOD1 polyubiquitination with primarily K6-, K27-, and K29-linkages. Note that no polyubiquitinated mutant SOD1 was recovered in the 4% stacking gel (panel A) and therefore removed prior to transfer in subsequent experiments. Mock refers to transfection with an equivalent amount of empty vector. Whole cell lysates (WCL) were loaded to demonstrate plasmid expression. Actin serves as loading control. Data is representative of 3 independent experiments.

Next, to determine the fate of mutant SOD1 with respect to TRAF6-mediated polyubiquitination, we performed ubiquitination assays with single-lysine ubiquitin mutants to elucidate the polyubiquitin chain linkages with which mutant SOD1 is modified (**Fig. 5B**). Single-lysine ubiquitin mutants have only one of seven lysines available (all other lysines are mutated to arginine) and thus can only be used to catalyze polyubiquitin chains linked through one particular lysine residue. We performed these assays with Flag-SOD1^A4V^ as we determined that this mutant had the strongest interaction with TRAF6 and was the most ubiquitinated. Our data suggest that TRAF6 promotes the polyubiquitination of mutant SOD1, with K6-, K27-, and K29-linkages prevailing (**Fig. 5B**, lanes 3, 4 and 6).

### TRAF6 is a modifier of mutant SOD1 aggregation

To address whether mutant SOD1 aggregation could be causally linked to the interaction with TRAF6, we performed filter retardation assays using 293FT cells expressing Flag-SOD1 (wild type or mutants), either following siRNA-mediated knockdown of TRAF6 (**Fig. 6A**) or co-expression of Myc-TRAF6 (wild type, ΔN, or C70A) (**Fig. 6C, E**). Cell lysates were filtered through cellulose acetate membrane filters, which trap protein aggregates with a diameter ≥200nm, and immunoblotted for Flag-SOD1. Western blots of cell lysates were included to verify the efficiency of TRAF6 knockdown, to demonstrate equal plasmid expression across conditions, and to provide an indirect loading control for the filter membranes (**Fig. 6B, D, F**). Note, cell viability was observed to be equivalent across conditions. Flag-SOD1 variants tested in the assays were: SOD1^A4V^ (high aggregation propensity and strong TRAF6 interaction), SOD1^G93A^ and SOD1^E100G^ (high aggregation propensity and less strong TRAF6 interaction). In experiments with TRAF6 knockdown, all SOD1 variants aggregated less in cells treated with two independent siRNAs for TRAF6 compared to siControl-treated cells (**Fig. 6A**, compare rows 2 and 3 to row 1). TRAF6 protein levels were effectively reduced with both siRNAs (**Fig. 6B**). In contrast, co-expression of TRAF6^WT^ or TRAF6^ΔN^, stimulated the aggregation of mutant SOD1 (**Fig. 6C**, columns 2-4 compare row 1 to rows 2 and 3), but not SOD1^WT^ (**Fig. 6C**, column 5). Taken together, these data indicate that lowering TRAF6 levels alleviates mutant SOD1 aggregation while the interaction of mutant SOD1 with the TRAF6 C-terminus is sufficient to exacerbate mutant SOD1 aggregation. This is consistent with the non-degradative ubiquitin linkage types observed.

**Figure 6:**
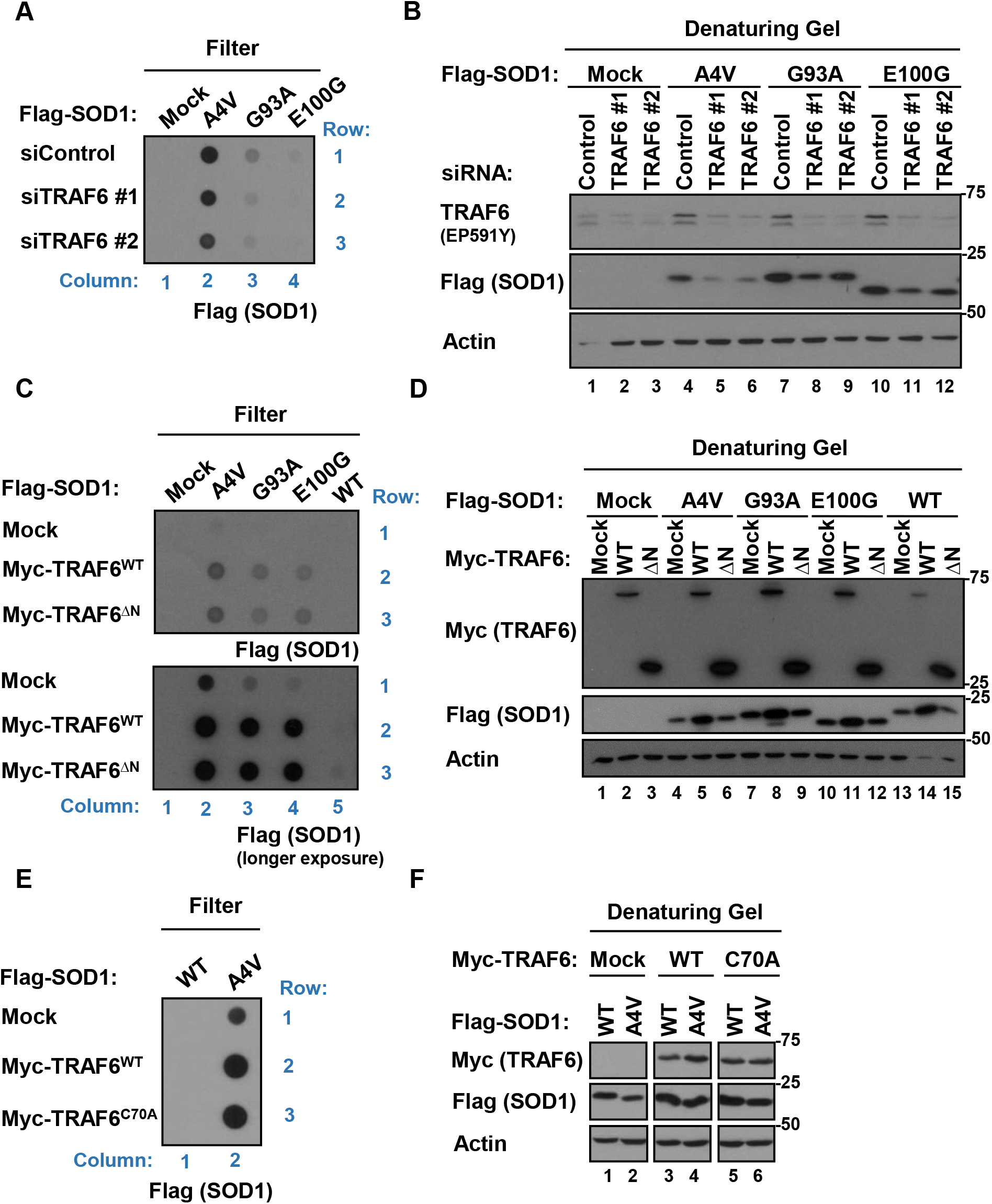
TRAF6 modifies mutant SOD1 aggregation but independently of its ubiquitin ligase activity. **(A, C, E)** Filter retardation assays in which protein aggregates ≥200nm diameter are captured on cellulose acetate membrane filters. Flag-SOD1 (wild type or mutants) was either co-expressed with Myc-TRAF6^WT^, TRAF6^ΔN^, or ubiquitin ligase-inactive TRAF6^C70A^, or Flag-SOD1 variants were expressed in 293FT cells that received siTRAF6 or siControl. Filter membranes were immunoblotted for Flag-SOD1. **(B, D, F)** Western blot of cell lysates to demonstrate equal plasmid expression and to verify TRAF6 knockdown across conditions. Immunoblots also serve as loading controls for the filter membranes. Mock refers to transfection with an equivalent amount of empty vector. Actin serves as loading control. All data is representative of 3 independent experiments.

Our results with the TRAF6 N-terminus harboring the RING domain, which is required for ubiquitin ligase activity, suggested that E3 ligase activity may not be implicated in SOD1 aggregation. To directly test this, we co-expressed mutant SOD1 with ubiquitin ligase-inactive TRAF6^C70A^. TRAF6^C70A^ stimulated SOD1^A4V^ aggregation comparable to TRAF6^WT^ (**Fig. 6E**, columns 2-4 compare row 2 to row 3), confirming that TRAF6 ubiquitin ligase activity is indeed not required for TRAF6-mediated mutant SOD1 aggregation. A similar result was obtained for SOD1^G37R^ (data not shown). Lastly, we examined whether TRAF6^C70A^-stimulated aggregation of mutant SOD1 correlates with a bimolecular interaction between TRAF6 and mutant SOD1. We observed that mutant SOD1 interacted comparably with TRAF6^C70A^ and TRAF6^WT^ (**Fig. S3**, compare lanes 7-8 to 10-11). Taken together, these data indicate that TRAF6 ubiquitin ligase activity is not required for TRAF6 interaction with mutant SOD1 nor TRAF6’s capacity to stimulate mutant SOD1 aggregation. We conclude that the interaction of mutant SOD1 with TRAF6 is sufficient to modulate mutant SOD1 aggregation.

### Mitochondrial recruitment of TRAF6 in SOD1^G93A^ rats

A pool of mutant SOD1 exists in an aggregated state on the surface of spinal cord mitochondria in SOD1^G85R^ mice and SOD1^G93A^ rats (18). In addition, we have demonstrated that B8H10 and AMF7-63-reactive misfolded SOD1 accumulate in mitochondrial fractions isolated from SOD1^G93A^ rat spinal cords and correlates with disease progression (10). We speculated that these observations could be linked to misfolded SOD1 interacting with TRAF6. TRAF6 is known as a cytoplasmic-localized protein, but its capacity to translocate to mitochondria in certain contexts has been previously reported (31,33,34). Since we detected TRAF6 as a binding partner of misfolded SOD1 at the mitochondria, we reasoned that TRAF6 must be recruited. To experimentally address this, we examined TRAF6 levels in spinal cord lysates and purified floated mitochondria from SOD1^G93A^ rats (**Fig. S4**). We collected samples at two different disease stages: 10 weeks (pre-symptomatic) and 14 weeks (pre-symptomatic/prior to disease onset). These two age groups were chosen for comparison because misfolded SOD1 is not readily detectable at spinal cord mitochondria in 10 week old SOD1^G93A^ rats but is at 14 weeks (10). Age-matched non-transgenic and SOD1^WT^-expressing rats were included as controls. Collectively, no differences in TRAF6 protein levels were observed in SOD1^G93A^ rat spinal cord lysates compared to controls and between age groups (**Fig. S4A**). Neither was there a difference observed in the level of TRAF6 in mitochondrial fractions compared to controls and between age groups (**Fig. S4B**). Surprisingly, however, we observed TRAF6 in mitochondrial fractions from non-transgenic animals, indicating that a pool of TRAF6 is constitutively localized to spinal cord mitochondria. These data suggest that misfolded SOD1 interacts with a mitochondrial-localized pool of TRAF6 rather than requiring TRAF6 to be recruited.

Lastly, we noticed that the TRAF6 (EP591Y) antibody we used to detect TRAF6 at 60kDa, corresponding to its reported size, additionally detected two faster migrating bands at ~48kDa and ~35kDa (both unreported previously) enriched in spinal cord mitochondrial fractions compared to lysates (**Fig. S4, Fig. S5C**). Using a range of tools and approaches, we validated that all three detected bands are specific to this antibody (**Fig. S5**).

### Effect of mutant SOD1 on TRAF6-stimulated NF-κB activation

Since the constitutive localization of TRAF6 at the mitochondria is unreported, the function of TRAF6 as a resident mitochondrial protein is unclear. Thus, we further investigated the functional repercussions of mutant/misfolded SOD1 binding to TRAF6 using a proof-of-principle approach. TRAF6 plays a central role in cytoplasmic signal transduction from a multitude of immune receptors in response to stimuli (induced NF-κB signaling) but is also crucial for cellular homeostasis through maintaining constitutive NF-κB activity (basal NF-κB signaling) (59). Constitutive NF-κB activity in the mature CNS is restricted to neurons and is required for neuronal survival (60,61). Glial cells have very low levels of constitutive NF-κB activity, but activity is strongly induced in inflammatory contexts (62). An upregulation of NF-κB activity is seen at late stages of disease in mutant SOD1 transgenic mice and primarily in glial cells, which is associated with neuroinflammatory processes (63,64). Thus, we reasoned that mutant SOD1 binding to TRAF6 may have repercussions on TRAF6’s capacity to transduce signals in the NF-κB pathway. To experimentally address this, we performed dual NF-κB luciferase reporter assays in cells expressing Myc-TRAF6 and an escalating dose of Flag-SOD1^WT^ or Flag-SOD1^A4V^, together with a *Firefly* luciferase NF-κB reporter and a *Renilla* luciferase internal control (**Fig. S6**). The amount of transfected Myc-TRAF6 plasmid was previously determined by dose titration and was chosen to yield ~50% NF-κB activation relative to maximal TRAF6-stimulated activation, thus allowing for the detection of decreased or increased NF-κB activity. SOD1^A4V^ was chosen because it most strongly interacted with TRAF6 in other experiments and SOD1^WT^ served as TRAF6-non-interacting control. Flag-p65/RelA was incorporated as positive control. The sole expression of Myc-TRAF6 increased NF-κB activity by 7-fold relative to basal level (as expected), but mutant SOD1 did not have a significant effect on this across the entire titration range. This suggests that in this cellular context, mutant SOD1 binding does not adversely affect TRAF6 in activating NF-κB and that the signaling transduction cascade remained intact.

### TRAF6 is expressed in the cell types affected in ALS

We and others have previously demonstrated in mutant SOD1 transgenic rodent models that misfolded SOD1 is primarily detected in motor neurons at disease end-stage and rarely in glial cells (8,10,16,65–67). To assess whether TRAF6 is expressed in these cell types, we attempted to detect TRAF6 expression by immunohistochemistry in SOD1^G93A^ rat spinal cords. Unfortunately, none of the four commercial antibodies we tried gave specific labeling in rats, mice or in cultured cells. As an alternative, we immunoblotted whole cell lysates prepared from human iPSC-derived motor neurons (**Fig. 7A**, **Fig. S7**) and mouse primary cortical neurons and astrocytes (**Fig. 7B**). TRAF6 was detected in all cell types, but we observed a difference in the ratio of the 60kDa-TRAF6 and the 48kDa band in mouse cortical astrocytes compared to neurons. Densitometry revealed that 60kDa-TRAF6 was 3-fold more abundant in neurons than in astrocytes, while the 48kDa band was 0.7-fold less abundant in neurons than in astrocytes (**Fig. 7C**). Moreover, in neurons 60kDa-TRAF6 was 6.5-fold more abundant than the 48kDa band and 1.4-fold in astrocytes. These data indicate that TRAF6 is expressed in the cell types that primarily harbor misfolded SOD1 *in vivo*. Thus, we explored the interaction between misfolded SOD1 and TRAF6 in *post-mortem* tissues from SOD1 mutation-carriers affected by ALS. While misfolded SOD1 was detected in the motor cortex of three independent SOD1^A4V^ cases (data not shown), we were unable to detect TRAF6 in misfolded SOD1 immunoprecipitates, possibly owing to the low overall abundance of TRAF6 in CNS tissues (68).

**Figure 7:**
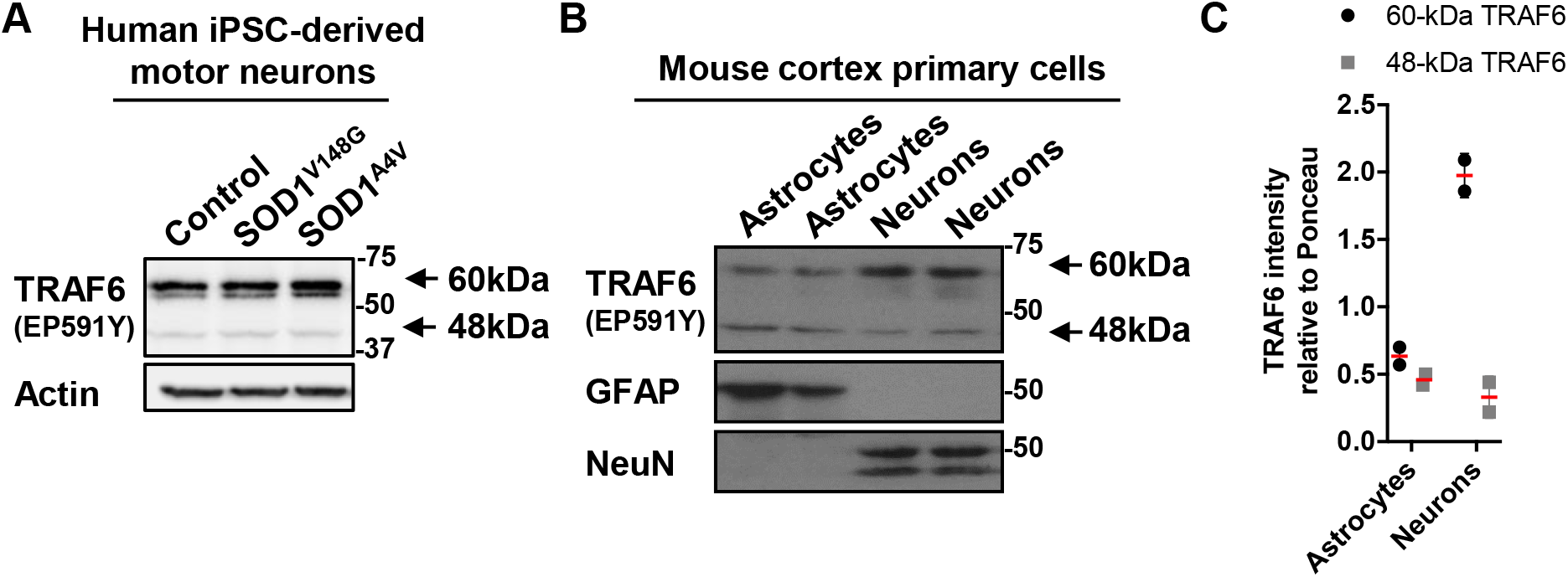
TRAF6 is expressed in ALS-disease relevant cell types. **(A)** Western blot of whole cell lysates from human iPSC-derived motor neurons differentiated for 14 days. Data is representative of 2 independent experiments. **(B, C)** Western blot of whole cell lysates from mouse primary cortical astrocytes and neurons and quantification of the abundance of TRAF6 relative to total protein stained with Ponceau S. Data represents the mean ± SD from two independent cell preparations. TRAF6 is detected in all cell types as a 60kDa immunoreactive band (corresponding to the reported size of TRAF6) and a faster-migrating band at about 48kDa (unreported).

## DISCUSSION

In this study, we conducted IP-MS proteomics using the SOD1^G93A^ rat model to identify the interactomes of misfolded SOD1 conformers at the mitochondria. Using antibodies reported to recognize distinct misfolded SOD1 conformers, we found a high degree of overlap for interactors (81%). Gene ontology analyses revealed a convergence on cellular responses related to unfolded proteins and protein ubiquitination. It is noteworthy that a number of interactors identified here are chaperone family members, which is in agreement with studies performed by others in different SOD1 models (69–72). In addition, we confirmed the previously reported interaction between human misfolded SOD1 and VDAC1 in rat spinal cord mitochondria (20). We also report that both misfolded SOD1 conformers interact with CCS, which may be expected given the known role of CCS in SOD1 maturation (73). However, the CCS-SOD1 heterodimer is an obligatorily transient complex (74). Thus, the detection of CCS in complex with misfolded SOD1 proteins here could reflect a sustained interaction that disrupts the available pool of CCS, thus impeding SOD1 maturation, as suggested by others (74). This could effectively contribute to a feed-forward loop of SOD1 misfolding. However, CCS has been previously ruled out as a modifier of the development of ALS-related phenotypes in transgenic SOD1 mice (75). Perhaps this concept should be revisited, given recent data in monozygotic siblings discordant for ALS where CCS is found to be down-regulated in the affected twin (76).

Interestingly, a few binding partners on our proteomics list were also selectively detected in B8H10 but not AMF7-63 IPs, and *vice versa* (**Table S2**). While the significance of these conformer-selective binding partners remains to be investigated, these data support the possibility that different misfolded conformers of SOD1 may have one or more distinct interactome-dependent pathomechanisms. One binding partner that was uniquely identified in B8H10 immunoprecipitates was 2-oxoglutarate dehydrogenase (*OGDH*) which has been described as an endogenous protein processed for mitochondrial antigen presentation (MitAP) in certain conditions (77,78). This raises the question as to whether misfolded SOD1 is itself presented on mitochondrial-derived vesicles (MDVs), similar to OGDH, or whether misfolded SOD1 may interfere with OGDH antigen presentation.

Another binding partner uniquely identified in B8H10 immunoprecipitates was ATP synthase subunit β (*ATP5B*), which has been previously reported as an interactor of apoSOD1 (metal co-factor deficient) in rat spinal cords (70). Consistent with this, B8H10 immunoprecipitates the metal-deficient mutants H46R, G85R and G127X comparably in our in cellulo experiments and we have previously shown that B8H10 preferentially recognizes apoSOD1 (10).

Interestingly, another ATP synthase subunit, ATP synthase F_0_ complex subunit β1 (*ATP5F1*), was uniquely recovered in AMF7-63 immunoprecipitates. Thus, the misfolded SOD1 conformers examined here each bound a different portion of the F_1_F_0_-ATP synthetase complex that drives mitochondrial ATP production. Defective mitochondrial ATP production has been previously reported in mutant SOD1-mediated disease and could positively contribute to compromised mitochondrial axonal trafficking (79).

We identified the E3 ubiquitin ligase TRAF6 as a novel binding partner of mitochondria-associated misfolded SOD1. We showed that SOD1 misfolding-associated exposure of the metal-binding loop and electrostatic loop in mutant SOD1 is a contributing factor for interaction with TRAF6. To this end, Garg *et al.* have performed *in silico* molecular dynamics simulations to further investigate the biophysical nature of the intermolecular interaction between mutant/misfolded SOD1 and TRAF6 (manuscript in preparation). It is our view, supported by this *in silico* work and our experimental evidence with SOD1^G127X^, that the electrostatic loop is not required for TRAF6 binding, since it is absent in this truncation mutant. This leads us to conclude that the interface for SOD1 binding to TRAF6 is formed primarily by residues located in the metal-binding loop, with a secondary contribution of residues in the electrostatic loop, as well as a potential co-option of the native SOD1 dimer interface. Interestingly, the conformational criterion for TRAF6 interaction is not universally fulfilled by all SOD1 variants. SOD1^V148I^, which is WT-like with respect to stability and structure (55–57), does not adopt a conformation that is characterized by the exposure of the B8H10 or AMF7-63 epitopes and does not interact with TRAF6 in cellulo. These data are reminiscent of the absence of an interaction of SOD1^V148I^ with Derlin-1, a functional component of ER-associated degradation (80). Indeed, all TRAF6-interacting SOD1 variants also interact with Derlin-1, suggesting that these mutants may collectively adopt a misfolded conformation that exposes a similar or possibly identical binding site for both interacting proteins. These findings certainly strengthen the notion that different SOD1 mutations may lead to motor neuron toxicity via distinct mechanisms. This possibility could be a contributor to the heterogeneity in disease onset observed for genetic subtypes of SOD1-ALS (81,82).

As previously shown in the context of other neurodegenerative diseases, TRAF6 stimulates the ubiquitination and aggregation of mutant/misfolded proteins (36,39,42,43). Here, TRAF6 stimulates the polyubiquitination of mutant SOD1 in an E3 ubiquitin ligase activity-dependent manner in cellulo. The polyubiquitin chain linkages we detected on mutant SOD1 were primarily K6, K27, and K29-linked, which is reminiscent of the type of chain linkages that have been found on other mutant/misfolded proteins when TRAF6 is co-expressed in cultured cells (36,39). TRAF6 also stimulated the aggregation of mutant SOD1, but surprisingly the TRAF6 N-terminus and E3 ubiquitin ligase activity were dispensable for this. Therefore, we conclude that these two events are independent, contrary to the currently proposed concept for the modification and autophagic degradation of mutant/misfolded protein aggregates (43). Moreover, polyubiquitinated mutant SOD1 was readily detectable in our assays in the absence of proteasome or autophagy inhibitors. This indicates that TRAF6-stimulated polyubiquitination and/or aggregation of mutant SOD1 does not prompt mutant SOD1 turnover but instead leads to its cellular accumulation. In fact, we observed in multiple experiments [e.g. **Fig. 2C** (2^nd^ blot from bottom, compare lanes 1 and 7), **Fig. 3A** (2^nd^ blot from bottom, compare lanes 1 and 4, 2 and 5), **Fig. 6D** (2^nd^ blot from bottom, compare lanes 13-15)] that the co-expression of TRAF6 resulted in increased mutant SOD1 levels in whole cell lysates. This was observed with TRAF6^WT^ and TRAF6^C70A^, but not TRAF6^ΔN^ or TRAF6^ΔC^, and was observed for TRAF6-interacting SOD1 variants as well as for TRAF6 non-interacting SOD1^WT^ and SOD1^V148I^. Collectively, this suggests that TRAF6 negatively affects mutant SOD1 turnover, but that this effect is not dependent on an intermolecular interaction between the two proteins and does not require TRAF6 E3 ligase activity. In line with this conclusion, we observed a reduction in the total level of mutant SOD1 in TRAF6-depleted cells (**Fig. 6B**), suggesting that lowering TRAF6 levels alleviates the aggregation of mutant SOD1 and prompts mutant SOD1 turnover. Taken together, this alludes to a scenario in which TRAF6 may act as a molecular scaffold serving a mechanical function in the context of mutant/misfolded SOD1 accumulation/aggregation at the mitochondria. That TRAF6 has E3 ubiquitin ligase-independent roles as a scaffolding protein has been previously demonstrated in the context of IL-1 signaling (83). However, given the role of TRAF6 in NF-κB signaling, which is intricately connected with regulating autophagy flux, the possibility remains that mutant SOD1 aggregation is indirectly and dynamically modulated in response to effects of TRAF6 on the autophagy machinery (28,29,84).

In addition to being localized to the cytoplasm, we found that a pool of TRAF6 is constitutively localized to mitochondria in the spinal cord. We propose that this mitochondrial TRAF6 interacts with and scaffolds the accumulation of misfolded SOD1. The TRAF6 (EP591Y) antibody we used for immunoblotting specifically detected TRAF6 at 60kDa in spinal cord mitochondrial fractions, and two other bands at 48kDa and 35kDa which were enriched in mitochondrial fractions compared to whole tissue lysates. This raises the possibility that there is more than one form of TRAF6 and that these forms may be differentially targeted to the cytoplasmic or mitochondrial compartments. How such short forms of TRAF6 are generated and what their function is at the mitochondria remains to be investigated.

TRAF6 is a multifaceted protein; it is a central player at the interface of inflammatory signaling and the autophagy machinery (27–30) and plays a role in mitophagy (34,35) and RNA metabolism (85,86), all of which are recurrent pathomechanistic themes in ALS, independent of genetic etiology. As such, we speculate that TRAF6 dysfunction due to mutant/misfolded SOD1 binding could affect cellular integrity at various levels, and in different cellular compartments. In fact, TRAF6 could participate in a variety of pathomechanisms underlying a broad spectrum of genetic subtypes of ALS, given that TRAF6 is reported to interact with many proteins already implicated in ALS (e.g. hnRNP A1 (86), Matrin-3 (86), Profilin-1 (86), Sqstm1/p62 (87), and TBK1 (88)). Our initial examination of the impact of mutant SOD1/TRAF6 on NF-κB signalling did not yield any differences. However, it is possible that the portion of TRAF6 bound to mutant SOD1 may be small compared to the total TRAF6 pool, which may explain our experimental outcome. Further investigations into the role of TRAF6 in ALS could yield important and unexpected insights which may help explain the heterogeneity and apparent complexity of the disease.

### Experimental Procedures

### Animals

Transgenic rats expressing human SOD1^WT^ and SOD1^G93A^ have been previously described (89,90). SOD1^G93A^ rats were bred and monitored biweekly as previously published (16). SOD1^G93A^ rats followed the expected disease course and reached end-stage at around 110 days. SOD1^G93A^ rats at 10 weeks and 14 weeks were pre-symptomatic, defined as no loss of body weight and intact hindlimb reflex. Animals described as symptomatic displayed bilateral hindlimb paralysis with no phenotypic involvement of the forelimbs. SOD1^WT^-expressing and non-transgenic control animals were age-matched. Animals of both sexes were used. Breeding, housing, and all manipulations were carried out in strict accordance with approved protocols from the CRCHUM Institutional Committee for the Protection of Animals (CIPA) and the Canadian Council on Animal Care (CCAC).

### Antibodies and siRNAs

Rabbit anti-AMF7-63 has been previously published (9,10). Commercially available primary antibodies were: mouse anti-Actin (MP Biomedicals, 69100), mouse anti-B8H10 (Medimabs, MM-0070), mouse anti-Flag (Sigma-Aldrich, F1804), rabbit anti-GAPDH (Cell Signaling, 5174), chicken anti-GFAP (Abcam, ab4674), rat anti-HA (Chromotek, 7c9), rabbit anti-Myc (Sigma-Aldrich, C3956), rabbit anti-NeuN (Novus Biologicals, NBP1-92693), rabbit anti-SOD1 (Enzo Life Sciences, ADI-SOD-101), rabbit anti-TRAF6 (EP591Y) (Abcam, ab33915, with ab183540 blocking peptide), mouse anti-VDAC1 (Millipore, MABNS04). Secondary horseradish peroxidase (HRP)-conjugated secondary antibodies were from Jackson ImmunoResearch: anti-chicken HRP (703-035-155), anti-mouse HRP (715-035-151), anti-rabbit HRP (711-035-152), anti-rat HRP (712-035-150). Stealth siRNAs were from ThermoFisher Scientific: siTRAF6 #HSS110968 (referred to as siTRAF6 #1), siTRAF6 #HSS110970 (referred to as siTRAF6 #2), and siRNA Negative Control #12935200 (referred to as siControl).

### Preparation of mitochondria and lysates from rat spinal cords

Rat spinal cords were homogenized in mitochondria isolation buffer I (10 mM Tris, pH 7.5, 1 mM EDTA, 210 mM D-mannitol, 70 mM D-sucrose) with protease inhibitors (10 μg/ml leupeptin, 10 μg/ml pepstatin A, 10 μg/ml chymotrypsin). Mitochondria were isolated by sedimentation and according to previously published protocols (91). For Fig. S4, spinal cords were homogenized in mitochondria isolation buffer II (20 mM HEPES-NaOH, pH 7.4, 1 mM EDTA, 250 mM D-sucrose) and mitochondria were isolated by buoyant density centrifugation on OptiPrep (Sigma, D1556) gradients as previously published (17). For tissue lysate preparations, an aliquot of homogenate was retained, adjusted to 1% v/v SDS and 1% v/v NP-40, and incubated for 10min on ice and 10min at ambient temperature. Extracts were mildly sonicated for 3 × 30 sec with 1 min icing between pulses with a Branson 2510R-DTH bath sonicator and cleared by centrifugation at 13,500*g* for 10 min at 4°C.

### Protein extraction, quantification and Western blot

Unless otherwise stated, whole cell lysates were prepared by extraction in RIPA buffer (50 mM Tris, pH 7.4, 150 mM NaCl, 1% v/v Triton X-100, 0.1% v/v SDS, 1% v/v sodium deoxycholate) with protease inhibitors (10 μg/ml leupeptin, 10 μg/ml pepstatin A, 10 μg/ml chymotrypsin) for 10 min on ice and 10 min at ambient temperature. Supernatants were recovered by centrifugation at 13,500*g* for 5 min at 4°C. Protein quantifications were performed with a BCA protein assay (ThermoFisher Scientific, 23227). Samples were boiled in 1x Laemmli buffer (60 mM Tris-HCl, pH 6.8, 2% w/v SDS, 10% v/v glycerol, 0.025% w/v bromophenol blue, 100 mM DTT) at 98°C for 5 min and separated on 12.5% Tris-glycine polyacrylamide gels. Gels were run at 100 V in running buffer (25 mM Tris, 192 mM glycine, 0.1% w/v SDS) and wet-transferred at 60 V for 1 hour at 4°C in transfer buffer (25 mM Tris, 190 mM glycine, 20% v/v methanol) onto 0.45 μm nitrocellulose membranes (Biorad, 1620155). Membranes were blocked in 5% milk-PBS/T (5% w/v Carnation^®^ instant skim milk powder in PBS, pH 7.4, 0.1% v/v Tween-20) for at least 30 min at ambient temperature. Primary antibodies were immunoblotted in 5% milk-PBS/T for 2 hours at ambient temperature or overnight at 4°C. For antibody pre-absorption with blocking peptide (Fig. S5), anti-TRAF6 (EP591Y) was diluted at 0.412 μg/ml in 5% milk/PBS-T, blocking peptide was added at 2-fold excess of the antibody and the solution was rotated end-over-end at ambient temperature for 1hr. HRP-coupled secondary antibodies were bound for 2 hours at ambient temperature. After primary and secondary antibody incubation, the membranes were washed three times with PBS/T. For all experiments, signals were captured on CL-XPosure™ radiography films (ThermoScientific, 34090) using ECL Western Blotting Substrate (Pierce, 32106); with the exception of Fig. 3A which was captured on the BioRad ChemiDoc system.

### Misfolded SOD1 co-immunoprecipitation from mitochondria and mass spectrometry

Rabbit anti-AMF7-63 (concentration 1.54 μg/μl) was coupled to Protein A Dynabeads (Invitrogen, 10002D) at full saturation according to bead binding capacity given by the manufacturer (6.15 μl per 40 μl beads). Mouse anti-B8H10 (concentration not provided by the manufacturer) was coupled to Protein G Dynabeads (Invitrogen, 10004D) at equivalent volume to AMF7-63. Beads coupled to isotype-matched ChromPure whole molecule IgGs (Jackson ImmunoResearch, 015-000-003 or 011-000-003) at full saturation were used as control. B8H10 and AMF7-63-coupled beads were crosslinked with BS^3^ (ThermoFisher Scientific, 21580), according to the manufacturer’s instructions. Beads were then tumbled with 150 μg spinal cord mitochondria isolated from symptomatic SOD1^G93A^ rats at 1 μg/μl final protein concentration in co-immunoprecipitation (IP) buffer (50 mM Tris-HCl, pH 7.4, 150 mM NaCl, 1 mM EDTA, 0.5% v/v NP-40) overnight at 4°C. The antigen-bound beads were washed three times with co-IP buffer and snap frozen in liquid nitrogen. For MS analysis, all samples were reduced on-bead with 5 mM DTT for 10 min at 95°C, alkylated with 5 mM iodoacetamide for 1hr at ambient temperature and trypsin-digested on-column for 18 hours at 37°C. Tryptic peptides were subjected to nanoLC-MS/MS analysis using a NanoAcquity UPLC (Waters) coupled to ESI-LTQ-XL-ETD mass spectrometer (ThermoFisher) as previously published (92). The .raw files were used to generate .mgf files using ProteoWizard (v3.0.18250) and submitted to the Matrix Science Mascot search engine (v2.6.2), to search against a rat Uniprot database (2019-02 release, # of entries 36,064) with human SOD1 sequence added (UniProt ID: P00441), and a decoy database (same database but randomized). Searches were performed with a specified trypsin enzymatic cleavage with one missed cleavage allowed and a 0.6 Da mass tolerance for precursor and fragment ions. Allowed modifications included carbamidomethyl at cysteines (+57 Da) as fixed and oxidation at methionines (+16 Da) as variable. Peptides showing Mascot scores ≥40 show an overall FDR less than 2.5% (decoy/target hits) and were determined as positive hits. The mass spectrometry proteomics data have been deposited to the ProteomeXchange Consortium (http://proteomecentral.proteomexchange.org) via the PRIDE partner repository with the dataset identifier < PXD015279> (93).

### Cell culture

HAP1 cells edited by CRISPR/Cas9 containing a 10 bp deletion in the first coding exon of TRAF6 (referred to as TRAF6-KO) and HAP1 parental cells (referred to as TRAF6-WT) (Horizon Discovery, HZGHC003466c011) were cultured in Iscove’s Modified Dulbecco’s Medium (ThermoFisher Scientific, 12440), supplemented with 10% fetal bovine serum (Wisent Bioproducts, 080150). 293FT cells (human embryonic kidney) were cultured in Dulbecco’s High Glucose Modified Eagles Medium (GE Healthcare, SH30081.01), supplemented with 10% fetal bovine serum and 2 mM L-Glutamine (Sigma, G7513). Primary mouse cortical neurons and astrocytes cultures were performed as previously described (94,95). For iPSC-derived motor neurons, motor neuron progenitors were generated from reprogrammed human fibroblast lines SOD1^A4V^ (Coriell, ND35671*C), SOD1^V148G^ (Coriell, ND35670*D), and healthy control line NCRM-1 (NIH) using a monolayer approach and a cocktail of small molecules optimized from published protocols (96). Motor neuron progenitors were then differentiated into motor neurons (50% DMEM/F12 and 50% Neurobasal medium, supplemented with N2, B27, L-Glutamax, antibiotic-antimycotic, 100 μM ascorbic acid, 0.5 μM retinoic acid, 0.1 μM purmorphamine, 0.1 μM Compound E, 10 ng/ml BDNF, 10 ng/ml CNTF, 10 ng/ml IGF-1) and cultured for 14 days. All cells were maintained at 37°C with a humidified atmosphere of 5% CO_2_. Studies with patient-derived cells abide by the Declaration of Helsinki principles and were conducted in accordance with approved protocols from the CRCHUM Institutional Review Board.

### Plasmids and cloning

pCMV-Myc-TRAF6^WT^ was a gift from Dr. Douglas Leaman (Wright State University, OH, USA) and has been previously described (97). pCMV-Myc-TRAF6^WT^ was used as a template to generate the deletion constructs TRAF6^ΔC^ (comprising aa. 1-288) and TRAF6^ΔN^ (comprising aa. 289-522) by classic restriction-based cloning into EcoRI and HindIII sites. pCMV-Myc-TRAF6^C70A^ was generated from pCMV-Myc-TRAF6^WT^ by site-directed mutagenesis with the QuikChange II Kit (Agilent, 200524) according to the manufacturer’s instructions. pCMV empty vector was used as a control. pcDNA3-Flag-SOD1^WT^, SOD1^G85R^ and SOD1^G93A^ were a gift from Dr. Jean-Pierre Julien (Université Laval, QC, Canada). pcDNA3-Flag-SOD1^WT^ was used as a template to generate pcDNA3-Flag-SOD1^A4V^, SOD1^G37R^, SOD1^H46R^, SOD1^E100G^, SOD1^V148G^ and SOD1^V148I^ by site-directed mutagenesis. pcDNA3-Flag-SOD1^G127X^ was subcloned from pCI-neo-SOD1^G127X^ (Gly127insTGGG), a gift from Dr. Don Cleveland (UCSD, CA, USA). pCI-Flag empty vector was used as control. pRK5-HA-Ubiquitin^WT^ and single-lysine mutants were a gift from Dr. Edward Fon (McGill University, QC, Canada) and are commercially available (Addgene, Ub^WT^ #17608, Ub^K6^ #22900, Ub^K11^ #22901, Ub^K27^ #22902, Ub^K29^ ^#^22903, Ub^K33^ #17607, Ub^K48^ #17605, Ub^K63^ #17606). pCMV-Flag-p65, pRL-null, and pGL3-P2(2X)TK for NF-κB luciferase assays have been previously published (98). All newly cloned plasmids were verified by Sanger sequencing (Genome Quebec). Cloning primers are listed in Table S5.

### Transfections

400 ng per expression plasmid, or respective empty vector (referred to as mock), were transfected into 293FT cells with Lipofectamine LTX with Plus reagent (Invitrogen, 15338030) according to the manufacturer’s instructions in Opti-MEM Reduced Serum Medium (ThermoFisher Scientific, 22600050). Normal growth medium was replaced after 3 hrs and cells were collected after 24 hrs. For siRNA delivery (150 pmol), 293FT cells were transfected with Lipofectamine 2000 (Invitrogen, 11668027) according to the manufacturer’s instructions. Normal growth medium was replaced after 5 hrs and cells were collected after 72 hrs. If cDNA expression plasmids were introduced into siRNA-treated cells, this was 24 hours prior to cell collection. For NF-κB luciferase assays, 293FT cells were co-transfected with pGL3-P2(2X)TK (25 ng, *Firefly* luciferase NF-κB reporter), pRL-null (50 ng, *Renilla* luciferase internal control), p65-Flag (0.5 ng, positive control), or Myc-TRAF6 (20 ng) and an escalating dose of Flag-SOD1^WT^ or Flag-SOD1^A4V^ (5-80 ng) with Lipofectamine LTX. Transfections were balanced with pCI-neo-Flag empty vector.

### In cellulo protein-protein interaction assays

293FT cells were transfected with the indicated plasmid combinations. Cell pellets were resuspended in PBS (137 mM NaCl, 2.7 mM KCl, 8 mM Na_2_HPO_4_, 1.5 mM KH_2_PO_4_, pH7.4) with protease inhibitors and lysates prepared by freeze-thaw extraction (3 cycles of freezing on dry ice and snap thawing in a 37°C water bath). The supernatants were recovered after centrifugation at 13,500*g* for 5 min at 4°C. Lysates were adjusted with 2x IP-buffer (100 mM Tris-HCl, pH 7.4, 300 mM NaCl, 2 mM EDTA, 1% v/v NP-40, with protease inhibitors) to 1.5x final concentration and tumbled overnight at 4°C with Dynabeads Protein G coupled to mouse anti-Flag or rabbit anti-Myc. Antigen-bound beads were washed three times with 1x IP-buffer and eluted by boiling at 98°C for 5 min in 2.5x Laemmli buffer. The eluates were analyzed by western blot as described.

### In cellulo ubiquitination assays

293FT cells were transfected with the indicated plasmid combinations. Whole cell lysates were prepared by extraction in RIPA buffer for 15 min on ice and were sonicated for 3 × 30 sec with 1 min icing between pulses with a Branson 2510R-DTH bath sonicator. Extracts were cleared by centrifugation at 13,500*g* for 10 min at 4°C. Dynabeads Protein G were coupled to rat anti-HA and 400 μg of cell lysate were immunoprecipitated as described.

### Filter retardation assays

293FT cells were transfected with the indicated plasmid combinations as described. Filter retardation assays were performed along previously published methods with slight modifications (99). Cells were lysed by freeze-thaw extraction in PBS with protease inhibitors. The DNA condensate was manually removed with a pipette tip and lysates were additionally digested with 0.35 KU DNaseI (Roche, 11284932001) per μl extraction buffer for 30 min at 37°C. Extracts were cleared by low-speed centrifugation at 800*g* for 10 min at 4°C. Supernatants were recovered, quantified and samples of 60 μg in a total volume of 150 μl PBS were loaded onto cellulose acetate membrane filters with 0.2 μm pore size (Whatman, 10404131) using the Bio-Dot^®^ Microfiltration apparatus (Biorad, 1706545). After 1 hr of passive filtration by gravity flow, the wells were vacated and washed three times with 200 μl PBS by gentle vacuum application. Filters were removed from the apparatus, blocked in 5% milk-PBS/T for 1 hr and immunoblotted as described.

### NF-κB luciferase assays

NF-κB luciferase assays were performed with the Dual-Luciferase^®^ Reporter Assay System (Promega, E1960) according to manufacturer’s instructions and as previously published (100). Briefly, 293FT cells were transfected with the indicated plasmid combinations in quadruplicate conditions. Luciferase luminescence was measured with a BioTek Synergy HT multi-mode microplate reader equipped with an automated substrate injection system. Data is expressed as the ratio of mean luminescence ± SD of *Firefly* luciferase/*Renilla* luciferase. For SDS-PAGE, quadruplicate samples were pooled and 30 μl were loaded.

### qRT-PCR

Human iPSC-derived motor neurons were differentiated for 14 days and total RNA was extracted with the miRNeasy Mini Kit (Qiagen, 217004) according to the manufacturer’s instructions. Reverse transcriptions were performed on 400 ng of total RNA extract in 40 μl total volume, using random primers (Invitrogen, 48190011) and M-MLV Reverse Transcriptase (Invitrogen, 28025013), according to the manufacturer’s instructions. Reactions were conducted in singleplex, in 10 μl total volume containing 1x Taqman Fast Advanced Master Mix (Applied Biosystems, 4444556), 1x Taqman primers/probe set (Thermo Fisher Scientific) and 1 μl of diluted cDNA. qRT-PCRs were performed on a QuantStudio 3 Real-Time PCR System (Thermo Fisher Scientific). Probes were from Applied Biosystems: ACTB (Hs01060665_g1), CHAT (Hs00758143_m1), GAPDH (Hs02786624_g1), HB9 (Hs00907365_m1), SOX1 (Hs01057642_s1). The data was analyzed using the deltaCT method (101).

### Software and Statistics

Enrichr was used to perform gene ontology (BP_2018, MF_2018, CC_2018) and pathway enrichment (KEGG_2019_Human) analyses (21,22). Protein sequence alignments were generated with MultAlin (102). Three-dimensional protein structure models were generated using iTasser (Iterative Threading Assembly Refinement) (103). The full-length model was constrained with PDB entry 3HCS.A, without alignment. No constraints were used for the deletion models. The highest C score models were chosen for display. Densitometry was performed in Adobe Photoshop CS4 and graphs were generated with GraphPad Prism 8. Statistics were performed in GraphPad Prism 8 (one-way ANOVA with Dunnett’s multiple comparisons or t-test, where appropriate).

## Acknowledgements

We thank Don Cleveland (UCSD), Heather Durham (McGill), Edward Fon (McGill), Jean-Pierre Julien (Laval), Douglas Leaman (Wright State), Giovanni Manfredi (Cornell), Diana Matheoud (U. Montreal), and Timothy Miller (Washington) for contributing reagents and/or for helpful discussions throughout the project. This work was supported by grants from the ALS Society of Canada (CVV), Muscular Dystrophy Association (CVV), Frick Foundation for ALS Research (CVV), and Rare Disease Foundation (SS/CVV). SS was supported by McGill University’s Faculty of Medicine, the Integrated Program in Neuroscience and the Centre de Recherche du CHUM. MG was supported by an FRQS Studentship. CVV is an FRQS Senior Research Scholar.

## Author Contributions

SS, MG, SP, CB, EH, and LD performed research. PG generated protein structure predictions, in collaboration with SSP. ATS and ASH conducted mass spectrometry analyses. YK cultured primary cells. Experiments on human iPSC-derived motor neurons were performed by MC, in collaboration with TMD (C-BIG iPSC-CRISPR platform). EC assisted with NF-κB luciferase assays, in collaboration with NG. NRC supplied the AMF7-63 antibody. AB, JFT, EMM, and JR provided advice. CVV, HMB, and SS conceived the study, designed experiments and interpreted the data. SS and CVV wrote the manuscript. All authors have reviewed and approved the manuscript prior to submission.

## The abbreviations used are

aa.: amino acids
ALS: amyotrophic lateral sclerosis
CC: coiled-coil motif
CSS: copper chaperone for SOD1
DSE: disease-specific epitope
fALS: familial ALS
GO: gene ontology
HC: IgG heavy chain
IP: immunoprecipitation
IP-MS: mass spectrometry-based immunoprecipitation proteomics
MATH: meprin and TRF homology domain
KEGG: Kyoto Encyclopedia of Genes and Genomes
LC: IgG light chain
NF-κB: nuclear factor kappa-B
RING: really interesting new gene domain
RS27A: ubiquitin-40S ribosomal protein S27A
sALS: sporadic ALS
SOD1: superoxide dismutase 1
TRAF6: TNF receptor-associated factor 6
RL40: ubiquitin-60S ribosomal protein L40
UBB: polyubiquitin-B
UBC: polyubiquitin-C
VDAC1: voltage-gated anion channel 1
WCL: whole cell lysate
zF: zinc finger motif

