## Supplemental Information for "TNF receptor associated factor 6 interacts with ALS-linked misfolded superoxide dismutase 1 and promotes aggregation"

**<u>Supporting Information:</u>**

<u>Supplemental Tables</u>
**Tables are provided as spreadsheets in separate file.*

Table S1*: Misfolded SOD1 IP-MS proteomics from SOD1$^{G93A}$ rat spinal cord mitochondria.

Table S2*: Misfolded SOD1 conformer-selective binding partners.

Table S3*: Significantly enriched terms in the GO sub-ontologies biological processes and molecular functions.

Table S4*: Significantly enriched KEGG pathways.

Table S5: Cloning and mutagenesis primers used for the generation of cDNA expression plasmids.

<u>Supplemental Figures</u>

Figure S1: Presence of ubiquitin in misfolded SOD1 immunoprecipitates from SOD1$^{G93A}$ rat spinal cord mitochondria.

Figure S2: Identification of TRAF6 in misfolded SOD1 immunoprecipitates from SOD1$^{G93A}$ rat spinal cord mitochondria detected by IP-MS proteomics.

Figure S3: TRAF6 E3 ubiquitin ligase activity is not required for interaction with mutant SOD1.

Figure S4: TRAF6 is not recruited to mitochondria with disease progression in SOD1$^{G93A}$ rats, but a pool of TRAF6 is constitutively localized to mitochondria.

Figure S5: Validation of the TRAF6 (EP591Y) antibody.

Figure S6: Mutant SOD1 does not modify TRAF6-stimulated NF-κB pathway activation.

Figure S7: Gene expression profile of human iPSC-derived motor neurons.

**Table S5: Cloning and mutagenesis primers used for the generation of cDNA expression plasmids.**

| **Mammalian Expression** | **Forward Primer**<br>**Reverse Primer** |
|---|---|
| **TRAF6$^{\Delta C}$** | 5’-GCGAATTCAGTCTGCTAAACTGTGAAAAC-3’<br>5’-CGAAGCTTCTAATACCCAGAGTCGGGTA-3’ |
| **TRAF6$^{\Delta N}$** | 5’-GCGCGAATTCATCTCAGAGGTCCGGAATTTC-3’<br>5’-CGAAGCTTCTATACCCCTGCATCAGTACT-3’ |
| **TRAF6$^{C70A}$** | 5’-CTGGAAAGCAAGTATGAAGCCCCCATCTGCTTGATGGC-3’<br>5’-GCCATCAAGCAGATGGGGGCTTCATACTTGCTTTCCAG-3’ |
| **SOD1$^{A4V}$** | 5’-GAATTCGCGACGAAGGTCGTGTGCGTGCTGAAG-3’<br>5’-CTTCAGCACGCACACGACCTTCGTCGCGAATTC-3’ |
| **SOD1$^{G37R}$** | 5’-GTGTGGGGAAGCATTAAAAGACTGACTGAAGGCCTG-3’<br>5’-CAGGCCTTCAGTCAGTCTTTTAATGCTTCCCCACAC-3’ |
| **SOD1$^{H46R}$** | 5’-GGCCTGCATGGATTCCGTGTTCATGA-3’<br>5’-CCAAACTCATGAACACGGAATCCATG-3’ |
| **SOD1$^{E100G}$** | 5’-GGCCGATGTGTCTATTGGAGATTCTGTGATCTCAC-3’<br>5’-GTGAGATCACAGAATCTCCAATAGACACATCGGCC-3’ |
| **SOD1$^{G127X}$** | 5’-GCATGAATTCGCGACGAAGGCCGTGTGC-3’<br>5’-GCTGCTCGAGTCATTTCCACCTTTGCCCACC-3’ |
| **SOD1$^{V148G}$** | 5’-CGTTTGGCTTGTGGTGGAATTGGGATCGCCC-3’<br>5’-GGGCGATCCCAATTCCACCACAAGCCAAACG-3’ |
| **SOD1$^{V148I}$** | 5’-CGTTTGGCTTGTGGTATAATTGGGATCGCCC-3’<br>5’-GGGCGATCCCAATTATACCACAAGCCAAACG-3’ |

**A**

| Identified proteins: | Ubiquitin | | | | | |
|---|---|---|---|---|---|---|
| Accession: | P62986 (RL40); P62982 (RS27A); P0CG51 (UBB); Q63429 (UBC) | | | | | |
| Peptides | Score (B8H10 IPs) | | | Score (IgG control) | | |
| | Replicate 1 | Replicate 2 | Replicate 3 | Replicate 1 | Replicate 2 | Replicate 3 |
| ESTLHLVLR | 48 | 50 | 45 | - | - | - |
| TITLEVEPSDTIENVK | 38 | 43 | - | - | - | - |
| TLSDYNIQK | 50 | 38 | 39 | - | - | - |
| Peptides | Score (AMF7-63 IPs) | | | Score (IgG control) | | |
| | Replicate 1 | Replicate 2 | Replicate 3 | Replicate 1 | Replicate 2 | Replicate 3 |
| ESTLHLVLR | 91 | 95 | 68 | - | - | - |
| TITLEVEPSDTIENVK | 109 | 52 | 90 | - | - | - |
| TLSDYNIQK | 104 | 61 | 60 | - | - | 35 |

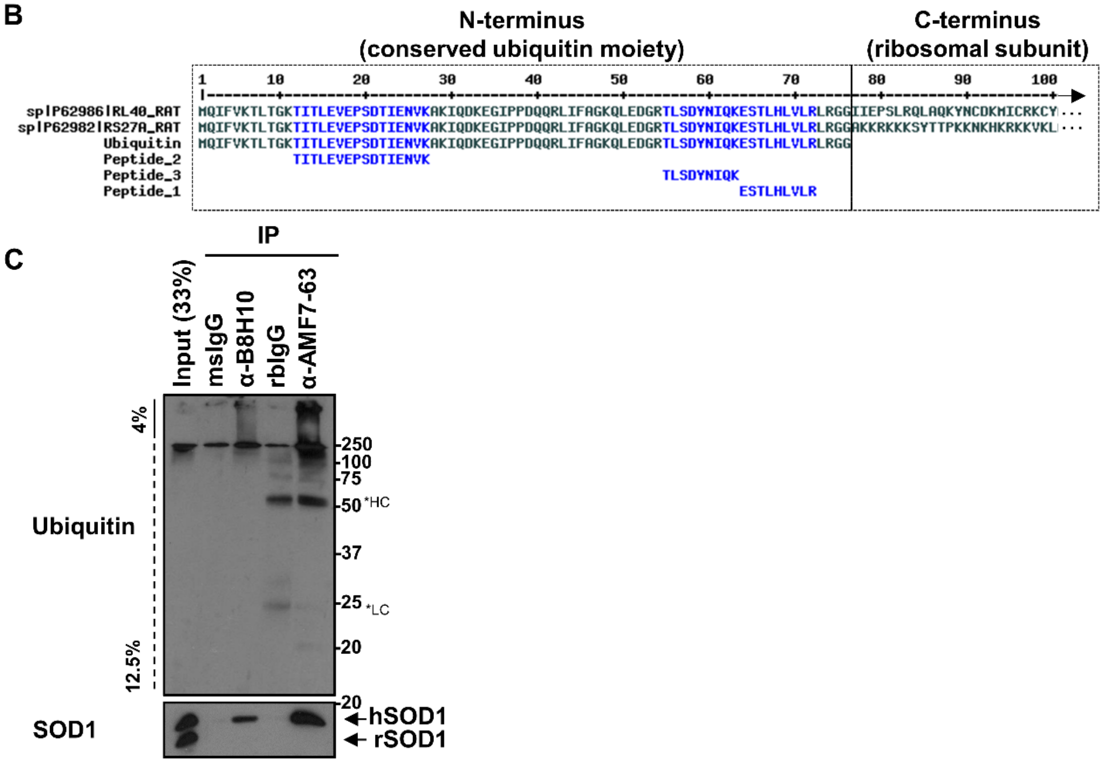


**Figure S1: Presence of ubiquitin in misfolded SOD1 immunoprecipitates from SOD1$^{G93A}$ rat spinal cord mitochondria. (A)** Ubiquitin was detected in B8H10 and AMF7-63 immunoprecipitates based on three peptides and was absent in IgG controls. **(B)** Alignment of the three peptides identified by IP-MS proteomics with ubiquitin (single moiety), RL40, and RS27A. All peptides align 100% with the N-terminal ubiquitin moiety of RL40 and RS27A. Note that these peptides would equally align with UBB and UBC polyubiquitin precursors, which are head-to-tail repeats of ubiquitin single moieties. **(C)** Detection of ubiquitin in B8H10 and AMF7-63 immunoprecipitates by western blot. HC, IgG heavy chain; LC, IgG light chain. Data is representative of 2 independent experiments.

**A**

| Identified protein: | TNF receptor-associated factor 6 | | | | | |
|---|---|---|---|---|---|---|
| Accession: | B5DF45 | | | | | |
| Sequence coverage: | 17% | | | | | |
| Peptides TRAF6 | Score (B8H10 IPs) | | | Score (IgG control) | | |
| | Replicate 1 | Replicate 2 | Replicate 3 | Replicate 1 | Replicate 2 | Replicate 3 |
| CEVSTRFDMGGLR | 84 | 99 | 85 | - | - | - |
| EIHDQSCPLANIICEYCGTILIR | 58 | 77 | 70 | - | - | - |
| GCRPEDPNYEETVK | 82 | 74 | 88 | - | - | - |
| LTILDQSEAVIR | 93 | 105 | 118 | - | - | - |
| METQSMHVSELK | 84 | 111 | 78 | - | - | - |
| QVSCVNCAVPMPYEEK | 69 | 56 | 84 | - | - | - |
| Peptides TRAF6 | Score (AMF7-63 IPs) | | | Score (IgG control) | | |
| | Replicate 1 | Replicate 2 | Replicate 3 | Replicate 1 | Replicate 2 | Replicate 3 |
| CEVSTRFDMGGLR | 46 | 41 | 38 | - | - | - |
| EIHDQSCPLANIICEYCGTILIR | 52 | 48 | 36 | - | - | - |
| GCRPEDPNYEETVK | 47 | 45 | 48 | - | - | - |
| LTILDQSEAVIR | 45 | 52 | - | - | - | - |
| METQSMHVSELK | 47 | 35 | 40 | - | - | - |
| QVSCVNCAVPMPYEEK | 41 | 41 | 42 | - | - | - |

**B**

>sp|B5DF45|TRAF6_RAT TNF receptor-associated factor 6 OS=Rattus norvegicus

10 20 30 40 50
MSLLNCENSC ASSQSSSDCC AAMANSCSAA MKDDSVSGCV STGNLSSSFM
60 70 80 90 100
EEIQGYDVEF DPPLESKYEC PICLMALREA VQTPCGHRFC KACITKSIRD
110 120 130 140 150
AGHKCPVDNE ILLENQLFPD NFAKREILSL TVKCPNKGCV QKMELRHLED
160 170 180 190 200
HQVHCEFALV ICPQCQRFFQ KCQINKHIIE DCPRR**QVSCV** **NCAVPMPYEE**
210 220 230 240 250
**KEIHDQSCPL** **ANIICEYCGT** **ILIR**EQMPNH YDLDCPTAPV PCTFSVFGCH
260 270 280 290 300
EKMQRNHLAR HLQENTQLHM RLLAQAVHNV NLSLRPCDAS SPSR**GCRPED**
310 320 330 340 350
**PNYEETVK**QL EGRLVRQDHQ IRELTAK**MET** **QSMHVSELK**R TIRSLEDKVA
360 370 380 390 400
EMEAQQCNGI YIWKIGNFGM HLKSQEEERP VVIHSPGFYT GRPGYKLCMR
410 420 430 440 450
LHLQLPTAQR CANYISLFVH TMQGEYDSHL PWPFQGTIR**L** **TILDQSEAVI**
460 470 480 490 500
**R**QNHEEVMDA KPELLAFQRP TIPRNPKGFG YVTFMHLEAL RQGTFIKDDT
510 520 530
LLVR**CEVSTR** **FDMGGLR**KEG FQPRSTDAGV

**C**

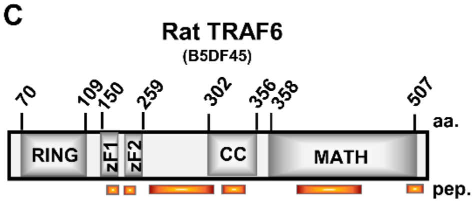


**Figure S2: Identification of TRAF6 in misfolded SOD1 immunoprecipitates from SOD1$^{G93A}$ rat spinal cord mitochondria detected by IP-MS proteomics. (A)** TRAF6 was detected based on six unique peptides in both B8H10 and AMF7-63 immunoprecipitates and absent in IgG controls. **(B, C)** The detected peptides (orange) cover 17% of the TRAF6 protein sequence. aa., amino acid; pep., peptide.

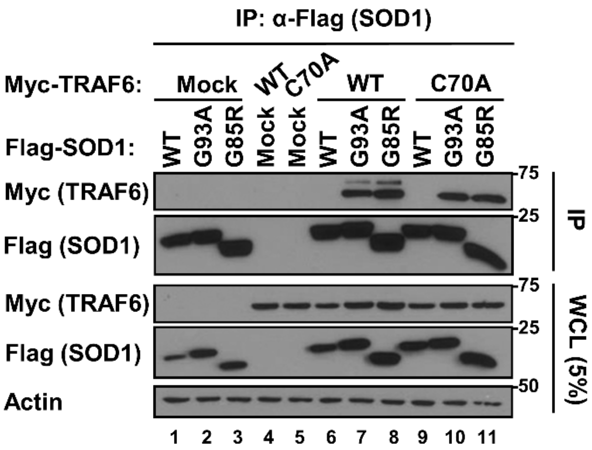


**Figure S3: TRAF6 E3 ubiquitin ligase activity is not required for interaction with mutant SOD1.** Flag-SOD1 (wild type or mutants) were co-expressed with Myc-TRAF6$^{WT}$ or the ubiquitin ligase-inactive mutant TRAF6$^{C70A}$ in 293FT cells. Co-immunoprecipitations were performed on Flag-SOD1 as bait and immunoprecipitates were analyzed for Myc-TRAF6 co-precipitation. Flag-SOD1 was blotted to demonstrate equal IP efficiency across conditions. Mock refers to transfection with an equivalent amount of empty vector. Whole cell lysates (WCL) were loaded to demonstrate equal plasmid expression. Actin serves as loading control. Data is representative of 3 independent experiments.

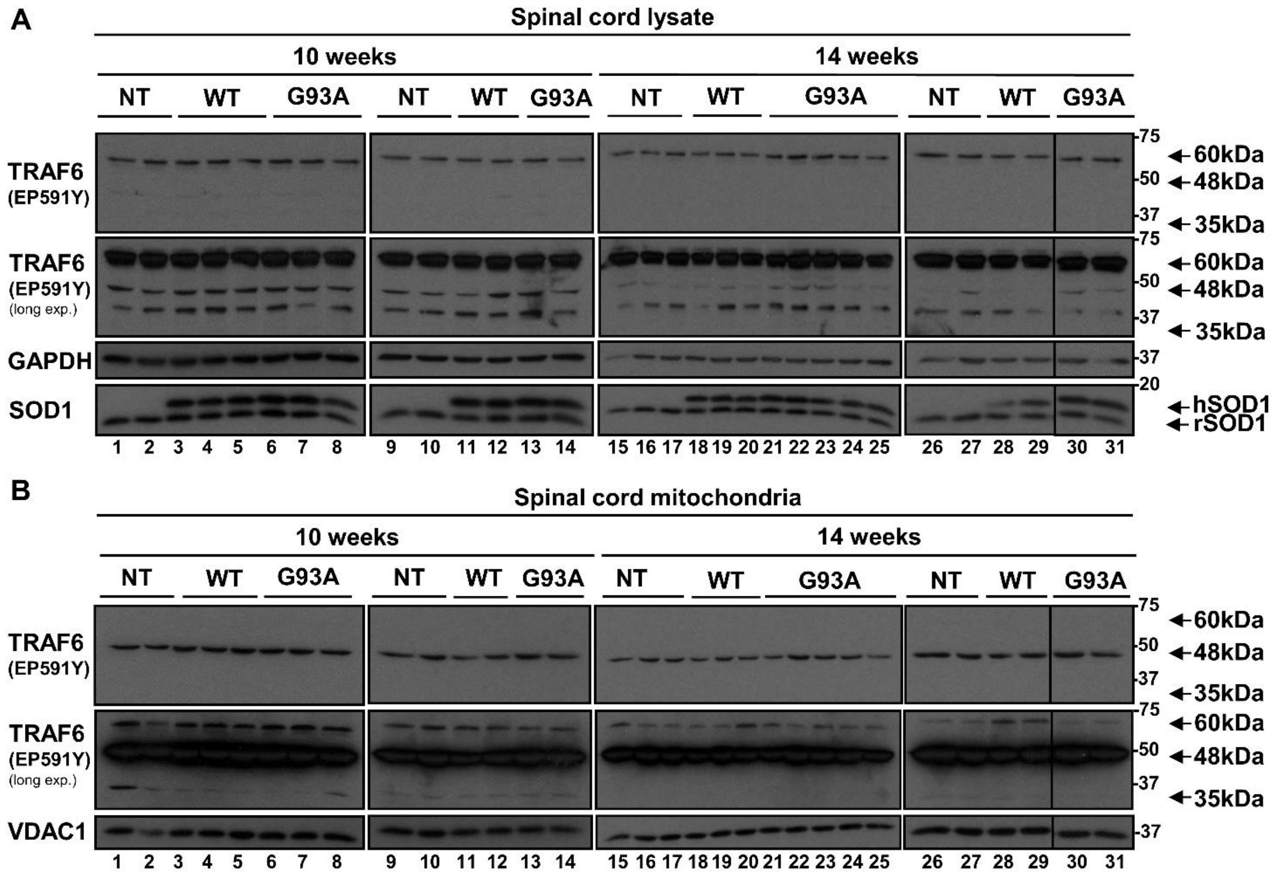


**Figure S4: TRAF6 is not recruited to mitochondria with disease progression in SOD1$^{G93A}$ rats, but a pool of TRAF6 is constitutively localized to mitochondria.** Western blots of **(A)** rat spinal cord lysates and **(B)** purified mitochondria from SOD1$^{G93A}$ rats at 10 weeks (pre-symptomatic) and 14 weeks (pre-symptomatic/prior to disease onset). Age-matched SOD1$^{WT}$ and non-transgenic rats served as controls. GAPDH and VDAC1 were used as loading control for lysates and mitochondrial fractions, respectively. SOD1 was blotted to show human and rat SOD1 expression in the transgenic animals.

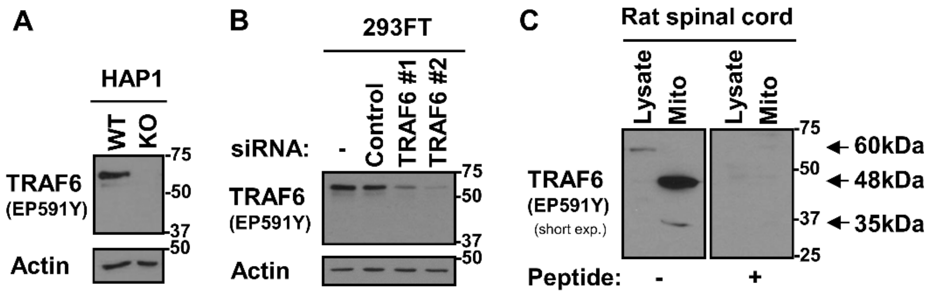


**Figure S5: Validation of the TRAF6 (EP591Y) antibody. (A)** Western blot of whole cell lysates from HAP1 TRAF6-WT and TRAF6-KO cells. EP591Y detects TRAF6 (60kDa immunoreactive band) in TRAF6-WT but not in TRAF6-KO cells. **(B)** Western blot of whole cell lysates from siRNA-treated 293FT cells. EP591Y detects changes in TRAF6 protein levels in siTRAF6-treated samples, but not in siControl-treated samples. **(C)** Western blot of rat spinal cord lysate and purified mitochondria with EP591Y, or EP591Y pre-adsorbed with its respective blocking peptide. 60kD, 48kDa and 35kDa immunoreactive bands were not detected with the pre-adsorbed antibody.

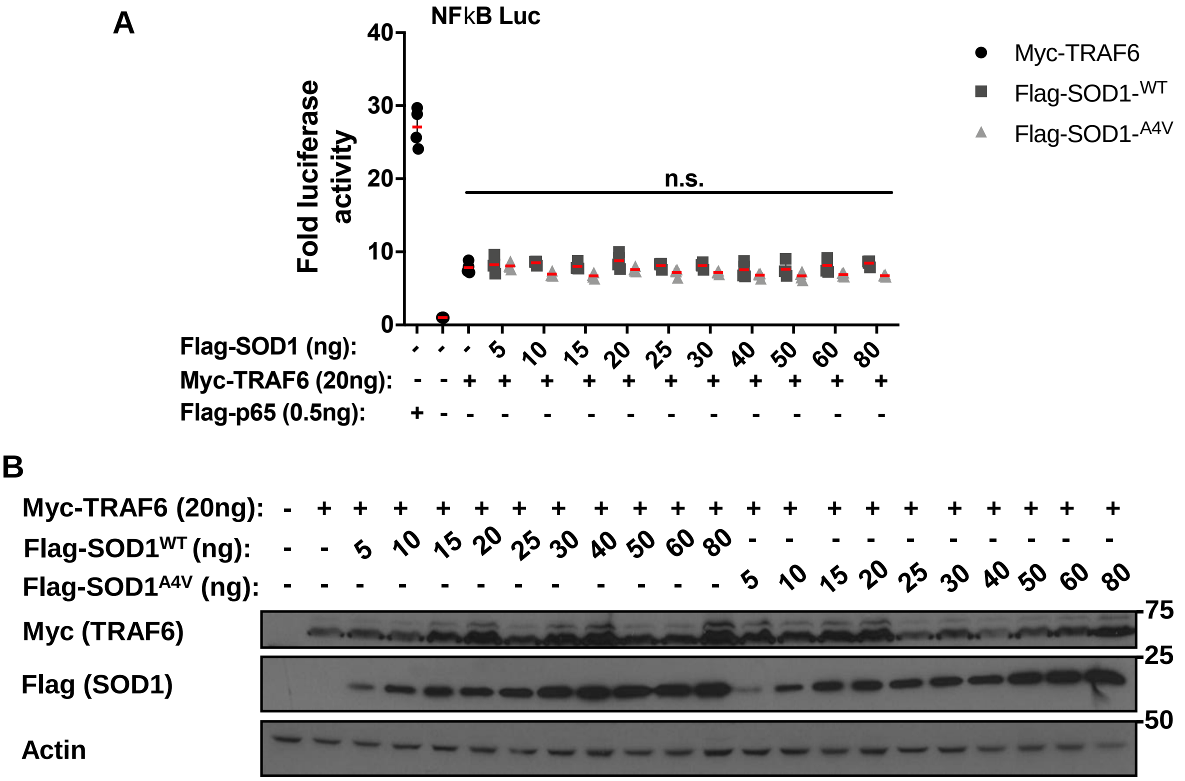


**Figure S6: Mutant SOD1 does not modify TRAF6-stimulated NF-κB pathway activation. (A)** Dual NF-κB luciferase reporter assay in 293FT cells transiently expressing a *Firefly* luciferase NF-κB reporter, a *Renilla* luciferase internal control, Myc-TRAF6, and Flag-SOD1$^{A4V}$ (TRAF6-interacting) or Flag-SOD1$^{WT}$ (TRAF6 non-interacting) at an escalating dose. The amount of transfected Myc-TRAF6 plasmid yielding ~50% NF-κB activation, relative to maximal TRAF6-stimulated activation, was previously determined by titration. Flag-p65/RelA was used as positive control. Mean luciferase activity, expressed as *Firefly*/*Renilla* ratio, was taken from 4 independently transfected replicates. Plotted mean ± SD values are relative to basal pathway activity (=mock transfection, second bar). n.s., not significant. **(B)** Western blot of the cell lysates to demonstrate equal expression of Myc-TRAF6 and expression of Flag-SOD1 across conditions. Actin serves as loading control.

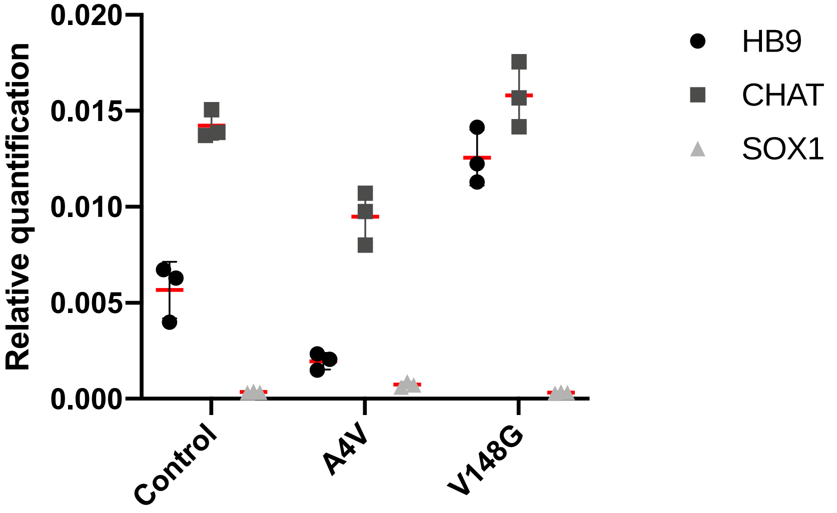


**Figure S7: Gene expression profile of human iPSC-derived motor neurons.** qRT-PCR on iPSC-derived motor neurons differentiated for 14 days are enriched for motor neuron markers HB9 and CHAT and not the neural progenitor marker SOX1. Plotted are relative quantification values (RQ = $2^{-\Delta CT}$) ± SD. Data was normalized to the mean of ACTB and GAPDH.
